# A high degree of clonal reproduction and aggregated structure of milkweed clones does not affect ramet sexual reproduction

**DOI:** 10.1101/2025.05.23.655745

**Authors:** H.M. Machiorlete, J.R. Puzey, H.J. Dalgleish

**Affiliations:** College of William & Mary, Department of Biology, 540 Landrum Dr, ISC 3050, Williamsburg, VA, USA

## Abstract

1. Clonal plants have unique population structures where genetic individuals can occupy many locations at once, but the effect of ramet spatial arrangement on demography is poorly understood.

2. We analyzed the spatial, genetic, and demographic characteristics of 798 ramets from four populations of *Asclepias syriaca*, a highly clonal plant, in northern Virginia.

3. *Asclepias syriaca* clones varied in size and spatial aggregation: ramets of the same genotype were more likely to be spatially clustered but when clones were small, we observed more spatial dispersion, potentially to avoid competition with self. Despite low genetic variation over all (Genotypes/Number of Individuals = 0.1), ramet sexual reproduction was not related to local genetic variation.

*Synthesis* High rates of clonality in *A. syriaca* spatially structures demographic and competitive dynamics without impeding sexual reproduction, highlighting how reproductive allocation shapes the population ecology of plants.

## Introduction

Clonal reproduction is ubiquitous in plants: nearly two thirds of angiosperms have the ability to clone, producing genetically-identical, physiologically-independent shoots (ramets) while also producing sexual offspring through flowering (Doust, 1981; Klimes et al., 1997; Herben & Klimesova, 2020). Reproductive allocation between sexual and clonal pathways affects the ecology, demography, and evolution of most plant species (Reekie & Bazzaz, 2005; Eilts et al., 2011; Li et al., 2018; Franklin et al., 2020). Despite the ubiquity of clonal reproduction across angiosperms, we have an insufficient understanding of clonality’s unique role in structuring these forces. Indeed, nearly 30% of plant population models ignore clonal reproduction altogether (Janovsky et al., 2017), presenting a gap that weakens our ability to understand the drivers of plant population change over time.

Clonal plants must balance their dual reproductive strategies to maximize fitness and minimize costs (Obeso, 2002; Vallejo-Marin et al., 2010). Sex is the only means of creating genetic variation for adaptive evolution. However, when pollinators (or genetically variable pollen) are unavailable, clonal reproduction ensures reproductive success while avoiding the accumulation of deleterious mutations from meiosis (Lanfear et al., 2013). Cloning allows for plants to quickly regrow following disturbance and compensate for tissue loss from herbivory (Pellissier et al., 2016; Evans et al., 2023). If clones are physiologically integrated, they can share resources among ramets, allowing for colonization of more microsites and higher resource acquisition (Steufer et al., 1994). Prolonged lifespans through clonal growth (increased generation time, Janovsky et al., 2017) allow for more opportunities for sexual reproduction, alleviating some tradeoffs between sexual and clonal reproduction (de Witte & Stocklin, 2010). However, clonal growth may be detrimental in the long term through the loss of genetic diversity that enables resilience in variable environments (Vallejo-Marin et al., 2010).

Clonality also creates a spatial dynamic for genotypes: instead of a single genotype occupying one space, the genet has the potential to colonize many locations, and through plasticity, display multiple phenotypes (Fransen et al., 2001; Alpert & Simms, 2002). Fine-scale environmental variation can drive phenotypic differences even among genetically identical ramets (Ricono et al., 2020), especially in plants that evolved in heterogeneous environments (Baythavong, 2011). For example, variation in leaf morphology in *Alternanthera philoxeroides* was high among ramets, but less so when the ramets were physiologically integrated (Xu et al., 2012). Important demographic traits, such as size class and reproductive stage, can vary among ramets of the same genet as well, as shown within large clones of *Convallaria keisikei* (Araki et al., 2009). Furthermore, ramet demography is often driven by environmental rather than genetic variation; in clonal grasses, soil characteristics and not genotype was among the strongest predictors of ramet emergence and survival (Latimer & Jacobs, 2012).

Despite the relevance of spatial relationships to the ecology and evolution of clonal plants, few studies have investigated the spatial population genetics of clonal plants. Ramets of the same genet may compete with one another for light, pollinators or other resources. Depending on the movement capability of the clonal organ (roots, runners, and rhizomes can spread more easily than corms and bulbs, for example), ramets of one genet may cluster together (phalanx growth) or spread out throughout their population (guerrilla growth, Doust, 1981). Phalanx increases intra-genetic density, thus self-pollination events are common as pollinators are more likely to deposit pollen on the same genetic individual (Charpentier, 2001); self-pollination either increases inbreeding depression or, if the plant is self-incompatible, increases pollen loss (Case & Ashman, 2005). For example, *Calystegia collina* ramets in high density, low genetic diversity outcroppings produced fewer seeds than those in high diversity areas (Wolf et al., 2001, Van Druenen et al., 2015). Guerrilla growth forms are better equipped to forage for resources in heterogeneous landscapes, but physiological integration could be less common in guerrilla species (Franklin et al., 2020), because the larger distance between ramets creates more opportunities to break off from the maternal plant. Understanding how spatial variation interacts with genotypic diversity to shape demography and fitness is a crucial first step toward revealing the ecological and evolutionary significance of clonality.

Common milkweed (*Asclepias syriaca* L.) is an excellent model species to investigate the spatial population genetics of clonal plants. Common milkweed produces ramets from adventitious root buds, which can develop anywhere along its roots (Woodson, 1954; Stamm-Katovich et al., 1988). This likely enables milkweed to disperse ramets far from its progenitor, allowing genets to intermingle and mitigate within-genet competition. Milkweed is self-incompatible (Wyatt & Broyles, 1994), which limits inbreeding depression, but also makes milkweed vulnerable to pollen discounting; selection may favor individuals that opt for clonal reproduction to reduce pollen loss (Vallejo-Marin et al., 2010). Previous thought held that milkweed populations were composed of a few large clones consisting of many ramets (Wilbur, 1976) and that sexual reproduction was rare. However, recent evidence challenged this model of milkweed population structure and posits that sexual reproduction occurs frequently, supporting genetic diversity (Pellissier et al., 2016; Bakacsy & Bagi, 2020; Ricono et al., 2020). For example, Ricono et al. (2020) found that the number of genotypes was 30% less than the number of ramets sampled; thus, recruitment through sex outweighed recruitment by clonal growth. Furthermore, the largest clone found by Ricono et al. (2020) was comprised of only seven ramets, and over half of sampled ramets were singletons. However, their transect sampling methodology may have insufficiently sampled their populations. While transect sampling is good at capturing genetic variation, it is liable to miss ramets within genets and bias ratios of clonal and sexual reproduction towards the latter (Arnaud-Haond et al., 2007).

To resolve our understanding of clonal population structure and its role in driving ecology and demography, we aimed to characterize clonality in common milkweed in Northern Virginia populations with high sampling effort. In addition, we investigated the spatial patterns of genetic variation and ramet demography to infer the ecological significance of clonality in this species. To these aims, we asked the following research questions:

How many clones are there in these populations and what are their size distributions?
How are ramets spatially arranged within and among clones?
How does genetic diversity and clone spatial structure affect sexual reproduction?
Does ramet fitness covary with genotype or spatial location?

## Materials and Methods

### Study organism

Common milkweed (*Asclepias syriaca*) is a clonal perennial plant ranging from the Eastern US to southern Canada (De La Mater et al., 2018). Cloning occurs from adventitious root buds, in addition to flowering. Milkweed produces inflorescences of 50-100 self-incompatible flowers. These flowers are pollinated with sacs of pollen (pollinia), dispersed by insect pollinators, and eventually produce seed pods (containing an average of 210 seeds ± 57 SD, Dalgleish et al., 2024). Reproductive success in milkweed has been linked to ramet size, as well as herbivory from specialist insect herbivores; these herbivores have evolved to sequester milkweed defensive chemistry (cardenolides) to feed on leaf tissue. Herbivory is known to reduce flowering, seed production, and clonal sprouting (Dalgleish et al., 2024).

Milkweed demography has seasonal and perennial characteristics. In northern Virginia, ramets resprout and seedlings emerge around late May to early June, flower beginning mid-June, then produce pods in August and September. Following this, all above-ground tissue dies, and genet rootstock remains dormant until the next season. Therefore, ramets behave as an annual population, but genets are perennial. The lifespan of genets is unknown, but milkweed populations have been observed to extirpate in our study area after ∼10 years (Dalgleish, personal observation).

| Populatio<br>n | Latitude | Longitude | Habitat and<br>management | Population<br>size | # ramets<br>genotyped | %<br>genotyped |
| --- | --- | --- | --- | --- | --- | --- |
| Blandy<br>Meadow<br>(BM) | 39.062705 | -78.062108 | Mixed grass prairie,<br>burned every three<br>years | 569 | 387 | 0.78 |
| Research<br>Village | 39.062225 | -78.071851 | Grassy hill,<br>unmanaged | 203 | 113 | 0.56 |

### Study sites

Four common milkweed populations in northern Virginia were sampled in 2023 (Table 1). Blandy Meadow and Research Village are found in University of Virginia’s Blandy Experimental Farm, and Turner Pond and Sky Meadows were in Sky Meadows State Park. Populations were designated as ∼30m x 30m sections that included the highest density of ramets; population boundaries were established each year, as milkweed ramets do not emerge in the same place annually.

**Table 1.**
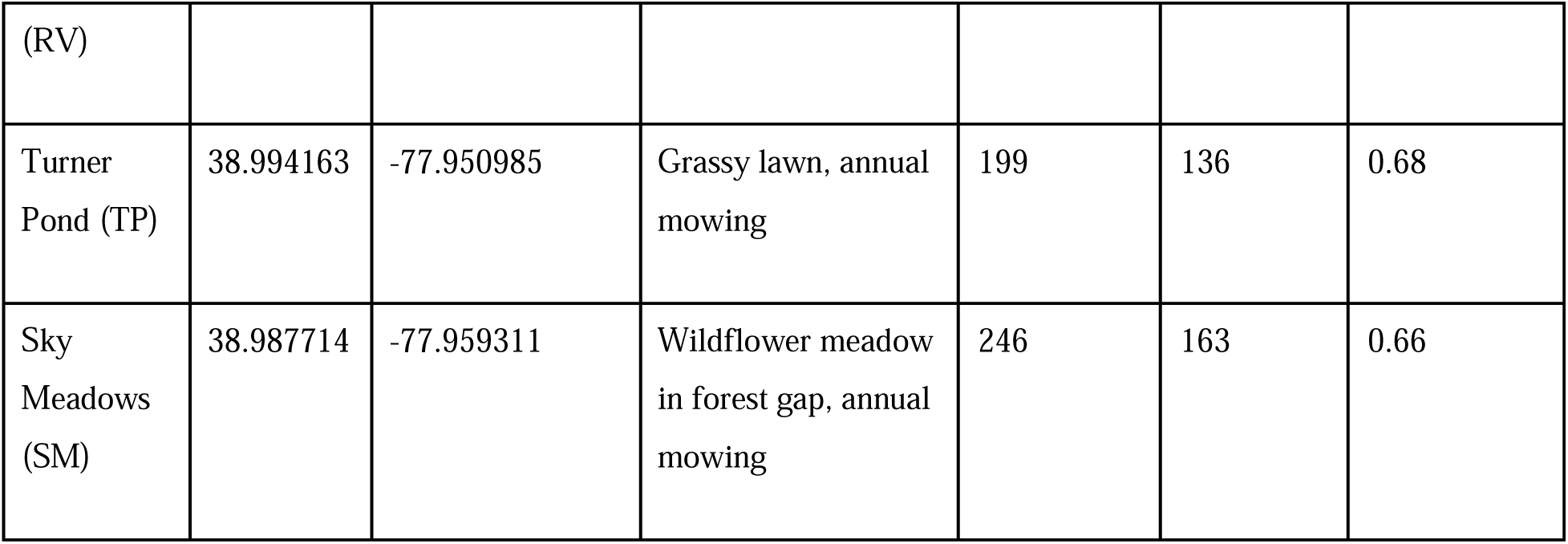
Description of study populations. Common milkweed population sizes estimated from the number of ramets mapped. Blandy Meadow and Research Village are found in Blandy Experimental Farm (University of Virginia) and Sky Meadows and Turner Pond in Sky Meadows State Park (Virginia Department of Conservation and Recreation).

### Field methods

#### Demographic data

For each ramet, we recorded data on ramet size and reproductive output. For ramet size, we measured apical height (stem base to the tip of the apical bud), stem width at the base, the length (tip to petiole) and width (at the widest section) of the largest leaf, and the total number of leaves. For reproductive output, we counted the number of inflorescences produced (hereafter called umbels) during peak flowering in late June, then returned in the late summer to quantify pod production. Pods were considered viable if they appeared large, green, and filled or large and brown, having released seeds previously; fruit structures that didn’t meet those conditions, or had stopped development, were counted as inviable pods.

We collected demographic data (size, survival, reproduction) in a population census. In mid-June, we flagged all individual ramets, then returned within 3 days to tag ramets and identify those who were missed on the first day; ramets that were missed over these two events were not included in the study. We recorded ramet size and flowering data in the same event, and herbivory and leaf data were collected separately within 3 days.

#### Spatial structure

To determine the spatial location of ramets, we mapped all ramets with the Postex Laser positioning system (Haglöf Sweden), where the position of a vertex was measured relative to transponders, providing relative coordinates with a 1 cm resolution. Ramets were mapped in mid-June following tagging and prior to demographic data collection.

#### Leaf sampling

We collected leaf samples (∼3cm x 3cm) from ramets in all populations. Leaf samples were stored in glassine bags and were transported in a cooler to a −80°C freezer for preservation within 12 hours of collection.

### Identifying clones

#### Selecting ramets for genotyping

To determine sampling scheme and intensity, we conducted a simulation study that varied the number of genets, their spatial structure, and sampling method (see Supporting Information for details). Sampling 75% of the population in randomly-selected clusters provided good estimates of clonal demography and spatial-genetic structure. Population sizes and the number of ramets genotyped are summarized in Table 1.

#### DNA extraction

Leaf samples from ramets selected for genetic analysis were extracted using Zymo Research Plant-Seed DNA Extraction 96-well Kit (Irvine, CA) with some modifications to the manufacturer protocol: Leaf tissue was flash frozen with liquid nitrogen and ground to a fine powder with a porcelain mortar and pestle, then approximately 80mg of tissue was transferred to a 2 mL microcentrifuge tube (to the 0.25 mL line). 400 µL of BashingBead Buffer and porcelain beads were added and tubes were vortexed for a minimum of 5 minutes. Next, tubes were centrifuged at 10,000 rcf for 2 minutes, and recoverable supernatant (∼200 µL) was transferred to 700 µL of Genomic Lysis Buffer with 0.5% v/v β-mercaptoethanol and thoroughly mixed. The Genomic Lysis Buffer mixture was centrifuged at 10,000 rcf for 2 minutes, then was transferred to a Zymo Research Silicon-A plate for wash and elution steps as detailed in the manufacturer’s protocol, with an additional dry spin (3,500 rcf for 5 minutes) following the second wash with g-DNA Wash Buffer.

We quantified DNA concentration and purity using a Nanodrop Spectrophotometer (ThermoFisher Scientific). DNA concentration ranged between 26 and 160 ng/µL and had high purity (A260/A280 > 1.8). A260/A230 ratios, corresponding to excess carbohydrates in eluted DNA, were consistently below 1.0, but DNA quality did not inhibit PCR or downstream processing.

#### Microsatellite markers

We amplified 7 microsatellite loci, which vary in the number of triallelic repeats, developed for common milkweed published in Kabat et al. (2010); we excluded the eighth locus, ASG5, because it consistently failed to amplify. All loci ranged in size between 87 and 434 base pairs (Table 2). The sizes we observed for ASB5 in our samples are different than published originally in Kabat et al. (2010) and Ricono et al. (2020).

**Table 2.** Microsatellite loci fragment sizes, allelic richness, and heterozygosity. Microsatellite loci were published in Kabat et al. (2010). Allelic richness, expected heterozygosity, and observed heterozygosity were calculated using *adegenet* (Jombart, 2008). Hardy-Weinberg equilibrium (HWE) tests were conducted from 1000 iterations of hw.test() in the *pegas* package (Paradis, 2010).

| Locus | Size range |  | Allelic Richness | Expected Heterozygosity | Observed heterozygosity | HWE X<sup>2</sup> | HWE df | HWE p |
| --- | --- | --- | --- | --- | --- | --- | --- | --- |
| ASC5 | 177 | 434 | 8 | 0.56 | 0.58 | 599.632 | 15 | <0.001 |
| ASF2 | 87 | 90 | 2 | 0.05 | 0.05 | 0.254 | 1 | 0.614 |
| AS94 | 143 | 167 | 12 | 0.73 | 0.82 | 1292.131 | 55 | <0.001 |
| ASB5 | 188 | 197 | 6 | 0.02 | 0.02 | 0.333 | 6 | 0.999 |
| ASH8 | 147 | 168 | 7 | 0.59 | 0.63 | 560.212 | 21 | <0.001 |
| ASF9 | 104 | 125 | 6 | 0.35 | 0.42 | 80.303 | 15 | <0.001 |
| ASG6 | 171 | 180 | 25 | 0.75 | 0.91 | 4638.608 | 171 | <0.001 |

#### PCR and fragment analysis

Multiplex PCR and fragment analysis were performed at the Cornell Institute for Biotechnology (Ithaca, NY), where all loci were amplified in a single reaction using Qiagen Multiplex PCR Master Mix under the following conditions: 2 µL template, 4 uM for all primers, and 1 µL bovine serum albumin. PCR reaction conditions performed as recommended using a 65°C annealing temperature. Primers were tagged with a GS-33 dye set (6FAM, NED, PET, VIC). PCR products were diluted 1:10, underwent the HiDi reaction, then were analyzed for fragment size on a 3730xl DNA Analyzer (Applied Biosystems) using a LIZ500 size standard.

#### Scoring microsatellites

Microsatellite alleles were scored using GeneMarker v. 3.2 (Soft Genetics). Fragment sizes were calibrated with the GS500 size standard (−250) setting; observed and expected fragment sizes of standards with scores less than 90 were removed from analysis. To classify alleles, we created an allele panel based on observed peaks from multiple populations. Alleles had to be present in at least two ramets to be considered a true allele. Due to fragment slippage during capillary electrophoresis (also known as stutter), few alleles had singular peaks. Therefore, stutter motifs that were consistent across samples were then classified into biallelic genotypes with the two most dominant peaks. For variants with two dominant peaks within 1 base of one another, we binned calls for either allele into one; of the two, we chose the call as the variant that was a multiple of 3. All alleles were checked by hand.

#### Population genetics

We investigated the population genetics of our sites and loci to check underlying assumptions of population structure and allele amplification success. First, to test for allelic dropout (the loss of alleles due to primer bias during PCR amplification), we calculated expected and observed heterozygosities for each locus. We tested whether all loci were in Hardy-Weinberg equilibrium (HWE) using the *hw.test* in the *pegas* package (Paradis, 2010) in R using 1000 iterations. Clonal plant populations are expected to violate HWE because cloning is a form of non-random mating resulting in a low effective population size that violates the infinite population size assumption (Arnaud-Haond et al., 2007; Douhovnikoff & Levanthal, 2016). We also checked for non-random allele associations (linkage disequilibrium, LD) with the standardized index of association (rd) using the *ia* function in *poppr* (Khamvar et al., 2014) with 1000 iterations; this would be expected in a clonal population, as cloning will perpetuate certain multi-locus genotypes, and thus allele combinations. Pairwise rd was also computed to investigate loci pairs in LD. Finally, to ensure that populations were independent of one another, we quantified population structure using pairwise F_st_ in *hierfstat* with 10,000 permutations (Goudet, 2004).

Using the same package, we calculated the inbreeding coefficient (F_is_, expected heterozygosity – observed heterozygosity / expected heterozygosity) for each population to check for reduced heterozygosity within populations.

#### Assigning ramets to genets

Prior to genet assignment, we generated a genotype accumulation curve using *adegenet* to ensure that we had sufficient loci to distinguish clones. We detected a plateauing effect, but did not have any redundant loci (Fig S1).

Microsatellite genotypes were combined into one multi-locus genotype to determine genet identities using the *allelematch* package in R (Galpern et al., 2012), consistent with previous research in this system (Ricono et al., 2020). This package compares repeat number similarity at each locus for each ramet-ramet comparison, where deviations are penalized and decrease the likelihood that they are a part of the same genet. These scores are aggregated and used to determine genet identity for each ramet. Because allelic diversity was low, multiple matches were common. In response, we did not allow any mismatches between samples and removed ramets that contained missing genotypes at any locus or loci. Therefore, our estimation of clonal membership is conservative.

### Data analysis

All analysis was performed in R v. 4.2.3 (R Core Team 2021).

#### How many clones are there in these populations and what are their size distributions?

We characterized the number of clones in all populations in 2023. Rates of clonal reproduction were described by the ratio of genotypes (G) to the number of individuals sampled (N), later referred to as G/N. We described the distribution of clone sizes as the number of ramets per genet, as well as the area occupied by clones. Clone area was quantified as the minimum convex polygon around ramets (using the *convexhull* function in the *spatstat* package; Baddeley et al., 2015). We used ANOVA to detect differences between sites in number of ramets per genet, clone area, and ramet density within clones.

#### How are ramets spatially arranged within and among clones?

For all sites, we modeled the probability of being a part of the same clone in response to inter-ramet Euclidean distance using a generalized linear model (GLM) with a binomial error distribution. In addition, we conducted Mantel correlations between genetic distance among ramets and inter-ramet distance within sites. Genetic distance was quantified as the Bray-Curtis distance among all alleles in all loci, which is shown to be an efficacious measure for populations with genetic structure (Shirk et al., 2017).

To understand growth patterns of clones, we quantified the arrangement of ramets within clones (those with 3 or more ramets). We simulated random distributions of ramets within clone area (*terra* package, 1000 iterations), then calculated and averaged the distance to the nearest neighbor of the same genotype for all ramets in the clone. We then determined whether the observed average nearest neighbor distance to self-ramets was significantly different from random (below the 0.05 quantile or above the 0.95 quantile of the mean). Ramets that were closer to ramets of the same clone than random had a clustered arrangement, and the opposite was considered dispersed. To assess whether clones of different sizes, and thus ages, employ different growth strategies, we performed an ANOVA on ramet growth pattern and the number of ramets per clone, as well as a Tukey HSD post-hoc test.

In addition, we aimed to understand how ramets of different genotypes interacted using two analyses. First, within the boundary of each clone, we counted the number of ramets belonging to that clone (self), and the number of ramets belonging to different clones (not self), to quantify intermingling (1-D, number of not-self ramets / all ramets in the clone area). Associations between intermingling and clone area were tested with a Spearman’s rank-sum correlation test.

Second, we calculated nearest neighbor distances between ramets of the same genotype and different genotypes and summarized the mean and variance of each category (self and not self) for all clones with more than 3 ramets. We compared means and variances of self vs. not-self using Wilcoxon tests.

#### How does genetic diversity and clone spatial structure affect ramet sexual reproduction?

Ramet sexual reproduction was measured as reproductive effort, success, and failure (or abortion rate). Reproductive effort was calculated as the number of umbels produced per cm of ramet height, reproductive success was the number of fertile pods produced per umbel, and reproductive failure was the proportion of pods that were aborted (see field methods for fertile pod classification).

To test the hypothesis that clonal intermingling reduced the likelihood of self-pollination, and increases fitness overall, we analyzed sexual reproduction of ramets using a clone- and a landscape-based analysis. Our clone-based analysis tested whether intermingling (1-D) was associated with reproductive failure.

Because intermingling within a clone is not spatially explicit, we aimed to search for covariance between fitness and genetic variation at the neighborhood scale with moving window analysis (MWA). First, we converted ramet points to rasters with a 10cm^2^ cell size using the *rasterize* function in *terra*. Then, MWA was conducted using *focal*. Circular window radii were 2m to match the distance at which there was a 50% chance of being the same clone. For each site, we quantified local G/N (number of genotypes relative to ramet density) and our three reproductive measures. Spatial autocorrelation of cell values was quantified using global Moran’s I from the *spdep* package. Following this, we correlated cell values for all pairs of response variables using Pearson’s correlations and adjusted p-values with Bonferroni corrections (to account for multiple comparisons). Correlation coefficients were then averaged across sites; pairings with coefficient confidence intervals that overlapped 0 were not significant.

#### Does ramet demography covary with genotype or spatial location?

To better understand the basis of ramet sexual reproduction, we examined the relationship between multivariate ramet sexual reproduction and clone, site, and subpopulation neighborhood. Neighborhood as opposed to x,y coordinates was used to account for patchiness within sites. Each site was subdivided into 16m^2^ neighborhoods; because sites were different sizes, the number of neighborhoods per site varied.

We used principal component analysis to compress variation in ramet sexual reproduction. PERMANOVA was used to test for differences among cluster centroids for our three groups: site, neighborhood, and genotype. In addition, we tested whether group variances were significantly different with permutation tests of multivariate homogeneity of group dispersion (performed with *beta.disper* and *permute.test* in the *vegan* package, Oksanen et al., 2024). These analyses were performed for all ramets in 2023, as well as within sites, where only neighborhood and genotype were compared.

To identify unique and shared explanations of variance in ramet phenotype, we performed redundancy analysis (RDA) using ramet sexual reproduction as a response matrix and genotype, neighborhood, and site as predictors. First, we tested that the model and predictor variables were significant using ANOVA (*anova.cca* in *vegan*). Finally, we used variance partitioning (*varpart* function) to deconstruct the variance explained by each response and discern the unique and shared explanatory power of each variable.

## Results

### Population genetic structure

Common milkweed populations in Northern Virginia were genetically distinct (Table 3). Pairwise F_st_ values were consistently above 0, and observed heterozygosity was either the same or higher than expected for all loci and populations (F_is_ was less than 0, Table 3). Therefore, we are confident that we can assess clonal identities with this data set. Many loci were in LD (rd = 0.25, p<0.001, Fig S2) and out of HWE, as expected with clonal species.

**Table 3.** F-statistics for four populations of common milkweed. 95% confidence intervals for population inbreeding coefficients (F*is*) and Nei’s pairwise F_st_ were bootstrapped using 10,000 iterations in *hierfstat*.

| Inbreeding coefficient ($F_{is}$) | | | |
| --- | --- | --- | --- |
|  |  | Lower 95% CI | Upper 95% CI |
| Blandy Meadow |  | -0.4342 | -0.0486 |
| Research Village |  | -0.3441 | -0.0459 |
| Sky Meadows |  | -0.6836 | -0.2927 |
| Turner Pond |  | -0.3463 | -0.042 |
| Fixation index (Pairwise $F_{st}$) | | | |
|  |  | Lower 95% CI | Upper 95% CI |
| Research Village | Blandy Meadow | 0.02705 | 0.20451 |
| Sky Meadows | Blandy Meadow | 0.20981 | 0.38516 |
| Turner Pond | Blandy Meadow | 0.06011 | 0.15336 |
| Sky Meadows | Research Village | 0.25653 | 0.46071 |
| Turner Pond | Research Village | 0.06988 | 0.15079 |
| Sky Meadows | Turner Pond | 0.13567 | 0.30156 |

### How many clones are there in these populations and what are their size distributions?

Common milkweed populations in northern Virginia were comprised of a low number of genetic individuals relative to the number of ramets sampled: we found only 81 clones out of 798 ramets sampled (G/N = 0.1, Fig 1, Table 4). Populations varied in their clonal diversity (Table 4). Sky Meadows exhibited the least amount of genetic variation, consisting of only seven clones among 163 ramets (G/N = 0.04); the population was dominated by a single clone comprised of 141 ramets. By contrast, Turner Pond had the most genetic diversity; its G/N was ∼0.23, between 2.2 and 5.3 times the genetic diversity of all other populations sampled.

**Figure 1.**
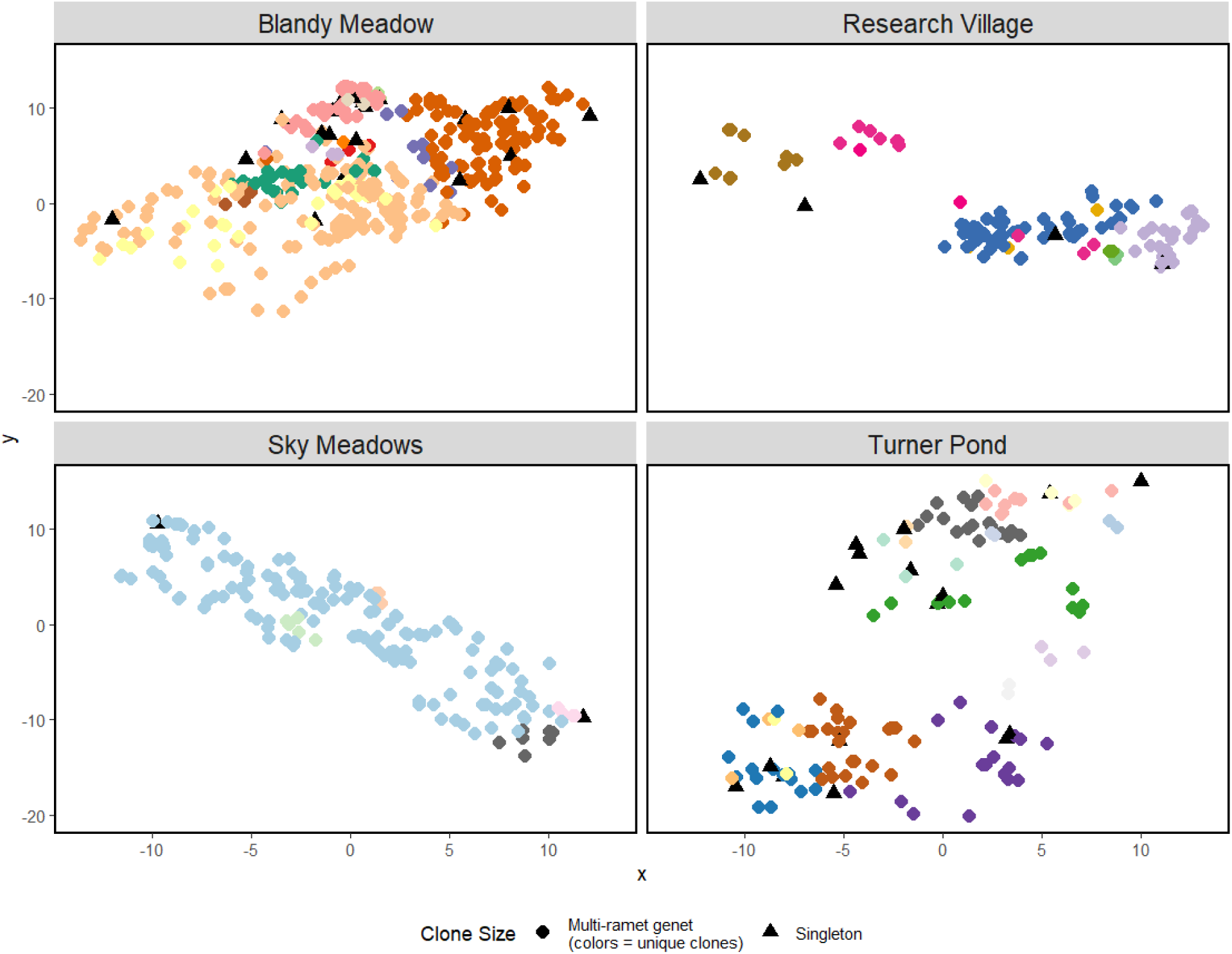
Common milkweed clones vary in size across populations. Clone identities (unique colors in each population) of milkweed ramets were determined for four milkweed populations in Northern Virginia in 2023. Clones were identified using 7 microsatellite loci (Kabat et al. 2010). Ramets were mapped to relative cartesian coordinates (x,y in meters) using the PosTex Mapping System (Haglof Sweden). Clones were not observed in multiple populations.

**Table 4.** High rates of clonal demography in common milkweed populations. We sampled up to 70% of ramets in four populations of common milkweed (N) and quantified the number of multilocus genotypes (G) using 7 microsatellite loci (Kabat et al. 2010).

| Site | # genotypes detected (G) | # ramets sampled (N) | G/N | # singletons | # multi-ramet clones | Largest clone size (# ramets) | Largest clone area (m<sup>2</sup>) |
| --- | --- | --- | --- | --- | --- | --- | --- |
| Blandy Meadow | 31 | 386 | 0.080 | 19 (61%) | 12 (39%) | 147 | 56.5 |
| Turner Pond | 31 | 136 | 0.228 | 16 (51%) | 15 (49%) | 23 | 39.6 |
| Sky Meadows | 7 | 163 | 0.043 | 2 (29%) | 5 (71%) | 141 | 66.3 |
| Research Village | 12 | 113 | 0.106 | 4 (33%) | 8 (67%) | 55 | 26.6 |

Among all populations, just over half (51%) of the 81 genets were singletons, and the remaining half were multi-ramet genets (Fig 1). Most of the multi-ramet genets had between 2 and 5 ramets (∼53%, 21/40), 25% of genets had between 5 and 20 ramets, 12.5% between 20 and 50, and the remaining 9.5% of genets had more than 50 ramets Though most genets were small, the largest clones composed as much as 86.5% of the ramet population (as in Sky Meadows), and as low as 16.5% of the ramet population (Turner Pond, Fig S3). Clones also varied in the amount of area they covered (on average 20.6 m^2^ ± 16.2 SD), as well as the density of ramets within their genet (1 ramet per m^2^ ± 0.68 SD on average), but clone area (ANOVA F_3,25_ = 0.38, *P* = 0.769) and ramet density within clones (F_3,25_ = 2.488, *P* = 0.0836) were similar across all four sites (Fig S4).

### How are ramets spatially arranged within and among clones?

Spatial-genetic structure was observed across all populations. Mantel correlations between Bray-Curtis genotypic distance and physical distance was significantly greater than 0 for all four populations in 2023 (Mantel correlation coefficients 0.2 – 0.41, p<0.05, Table S1). Therefore, ramets that were further apart were more genetically dissimilar.

Furthermore, genetic structuring was highly clustered (Figs 1, 2). Distance between ramets significantly predicted whether two ramets were from the same clone (Table S2 summarizes model fits). Within 1 meter, ramets in all populations had a high probability of being the same genetic individual, ranging between 0.55 (Blandy Meadow) to 0.85 (Sky Meadows). The likelihood of clonal membership significantly declined with increasing distance between ramets, but the rate of this decline varied across populations (Fig 2). Blandy Meadow, Turner Pond, and Research Village had similarly shaped probability curves; the inter-ramet distance at which the 50% probability mark (i.e. an equal probability of being the same or different genotype) was reached was at inter-ramet distances under 4 meters (1.8m for Turner Pond and Blandy Meadow, 4.1m for Research Village). However, for Sky Meadows, ramets within 29.9 meters were still more likely to be the same clone than not. With this model, ramets in Sky Meadows need to be over 100 meters away to have no chance of being the same clone.

**Figure 2.**
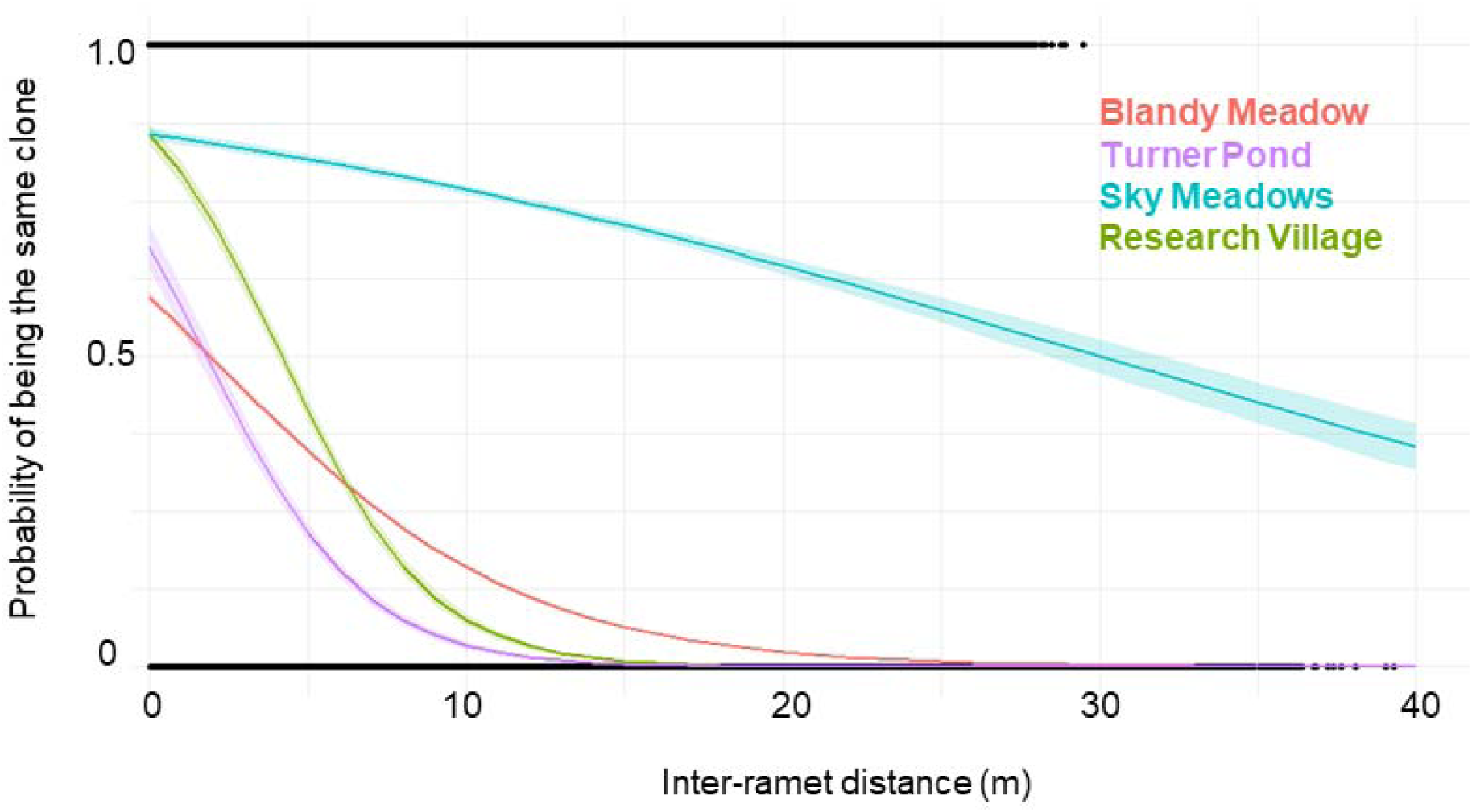
Clonal membership probability declines with increasing distance between ramets. Euclidean distance and genetic similarity (1-same, 0-different clone) were determined for ramets in four populations (see line colors) in 2023. Ramet-ramet comparisons (black points) were used to model the probability that two ramets belong to the same clone as a response of inter-ramet distance with a binomial generalized linear model. Lines represent model fits for each population ± 95% confidence intervals.

Within clones comprised of 3 or more ramets, ramet arrangement was variable. About half of clones (15 of the 29) had a dispersed arrangement of ramets meaning that their ramets were further than expected by chance to self-ramets than non-self-ramets. Only 4 of the 29 genets had a clustered arrangement (ramets were closer to self than non-self), and the remaining 10 were randomly distributed. Average nearest-neighbor distances to a ramet of the same genotype was the smallest for clustered clones (0.51 m ± 0.07 SD), and greatest for dispersed clones at 1.43m ± 1.04 SD meters, and in between for random (0.73 m ± 0.28 SD, Fig 3A). Ramet arrangement was associated with genet size (Fig 3B): genets with dispersed patterns had significantly fewer ramets (median of 4 ramets) than genets with random (24 ramets) or clustered (118.5 ramets) arrangements (*P* < 0.001 for each comparison).

**Figure 3.**
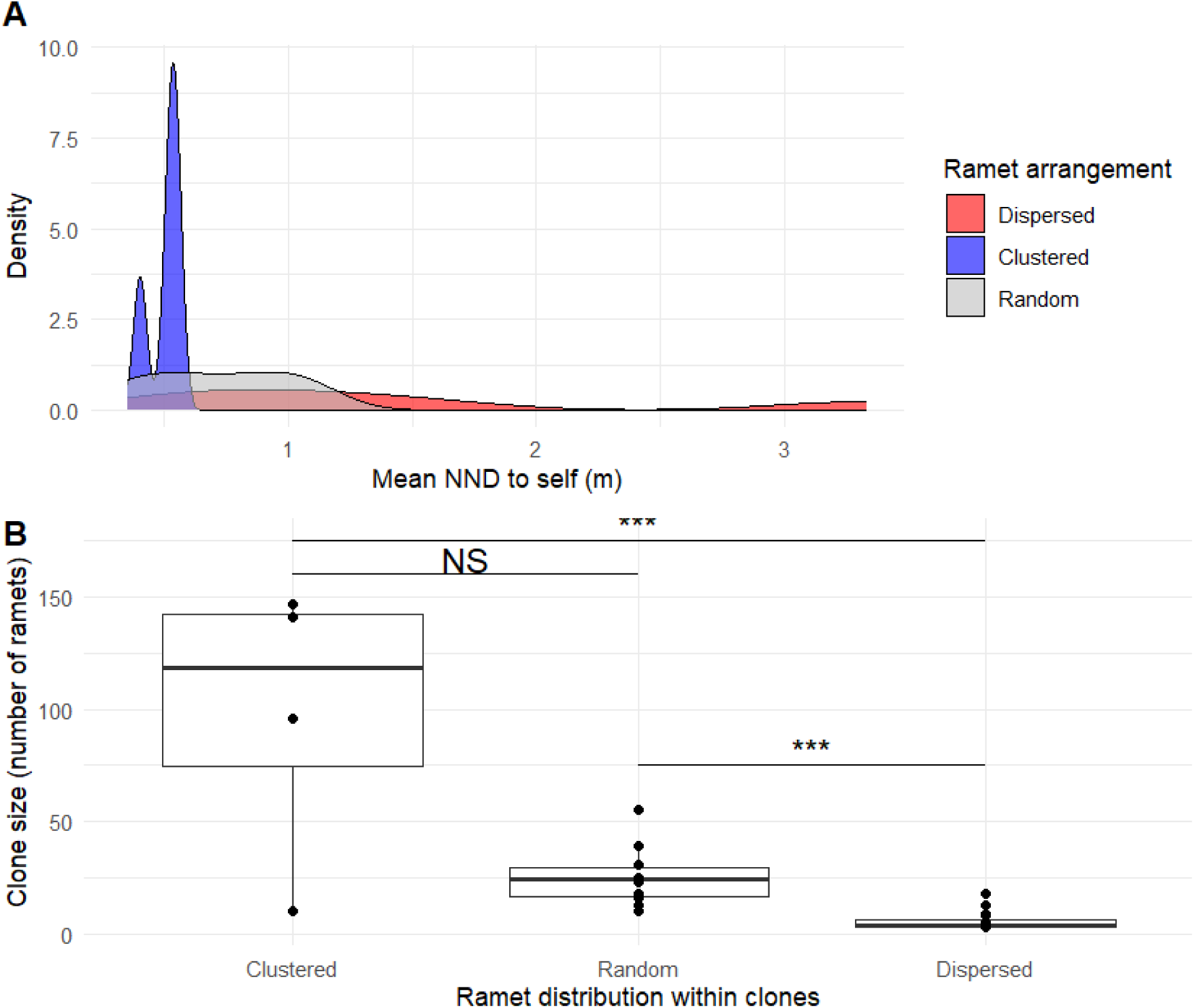
Ramet distributions within clones are related to clone size. Within the area of each clone (>2 ramets), we simulated a random distribution of ramets and calculated the nearest neighbor distance (NND) to self for all ramets within the clone (1000 simulations) and compared the observed mean NND to self with that of the random distribution (permutation test). Dispersed indicates a nearest neighbor distance significantly greater than random (above the 0.95 quantile of random), and clustered less than random (below the 0.05 quantile). A) Variation in mean NND to self for all clones by ramet arrangement. B) Relationship between ramet distribution and the clone size (the number of ramets per clone). Horizontal bars indicate post-hoc comparisons with a Tukey HSD test, *** = P<0.001, NS = not significant.

### How does genetic diversity and clone spatial structure affect ramet sexual reproduction?

Clone size influenced the degree of intermingling among genotypes: high intermingling, i.e. a higher proportion of non-self ramets within a clone’s area, was positively correlated with clone area (Spearman’s rho = 0.59, Fig 4). Therefore, as a clone colonized more area, more intermingling with other clones occurred.

**Figure 4.**
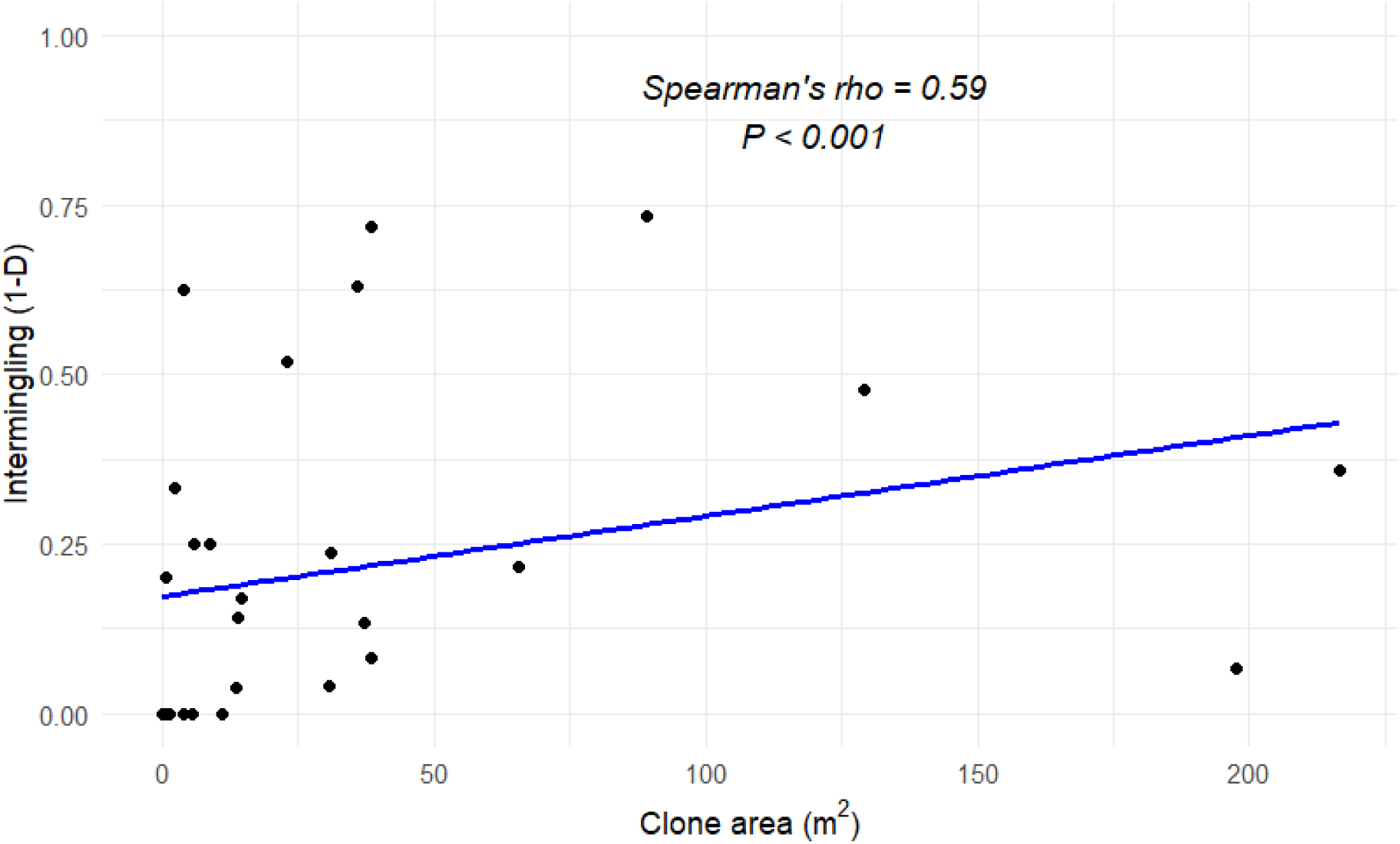
Clone intermingling increases with clone area. We generated a minimum convex polygon around all clones with 3 or more ramets and counted the number of ramets of the same genotype and different genotypes within the clone area. Spearman’s rank correlation was performed to determine associations between clone area and the proportion of ramets within a clone boundary belonging to different clones (1-D).

Reproductive success of ramets (number of pods produced per inflorescence) as well as reproductive failure (proportion of pods aborted) was related to clone area; reproductive success (rho = 0.41, S = 2376.8, *P* = 0.025) and failure (rho = 0.45, S = 2239.7, p-value = 0.015) increased with greater clone area. However, reproductive effort (number of inflorescences produces/cm height, *P* = 0.1) was not related to clone area. Ramet reproductive success was not related to clonal intermingling (*P* > 0.7). However, clonal intermingling did increase reproductive failure (the number of pods aborted/pods produced) per ramet (rho = 0.42, S = 5748, *P* = 0.025).

Moving window analyses for each population revealed the patchy distribution of genetic diversity (as measured by local G/N) and several measures of reproduction, identifying spatial hotspots for each (Figs 5, S5). In addition, we found high spatial autocorrelation for genetic diversity and all measures of reproduction (Moran’s I range = 0.77 – 0.97, Fig S6). However, local G/N was not significantly correlated with any of the reproductive measures (Fig 5), but all measures were positively correlated with one another, meaning that higher reproductive effort corresponded to higher pod production (rho = 0.52) and ovule abortion (rho = 0.51), including a positive association with reproductive success and failure (rho = 0.60). We show the results for Blandy Meadow, but spatial structure of ramet sexual reproduction was similar in all sites (see Fig S5, Table S3).

**Figure 5.**
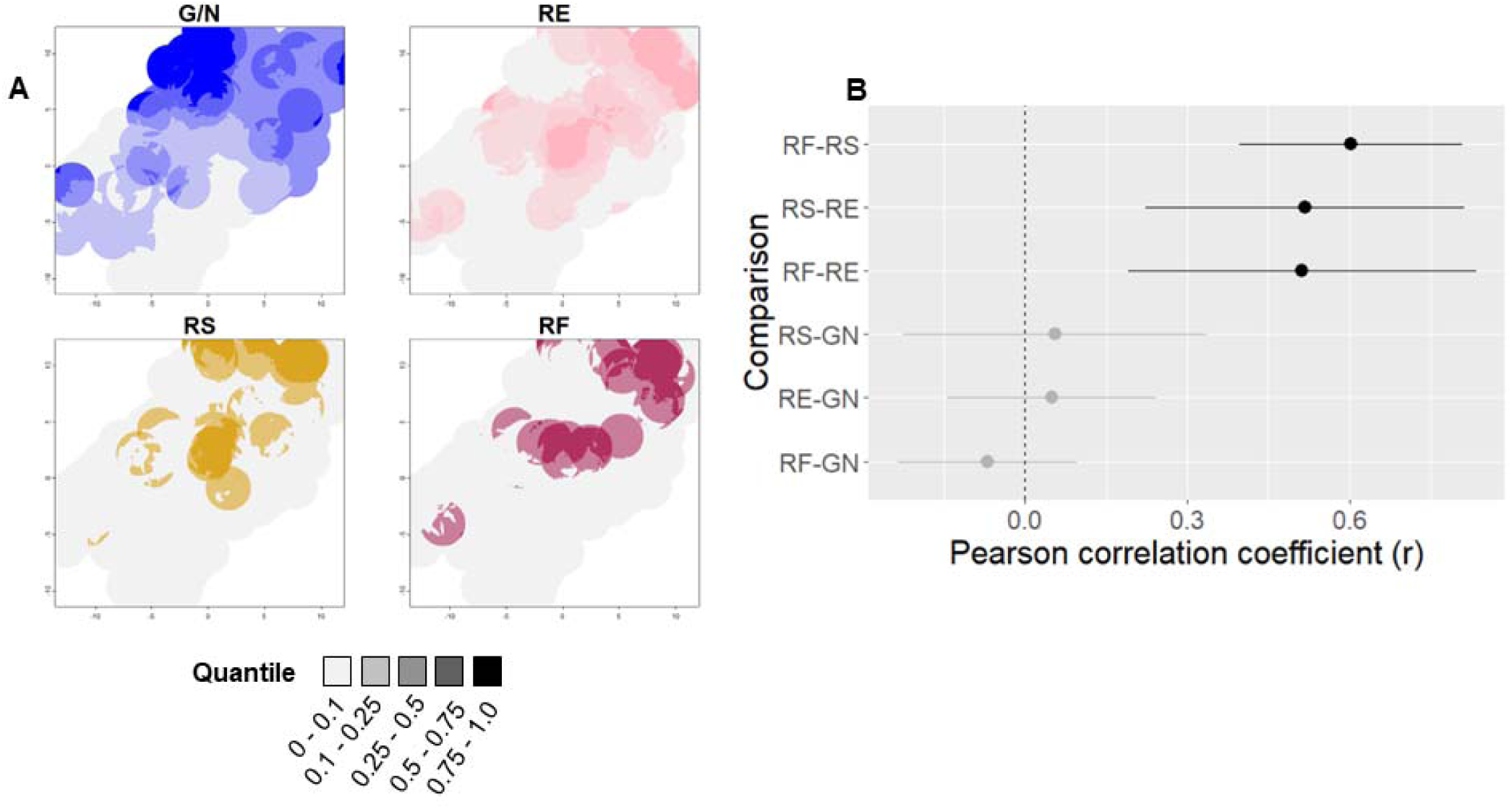
Weak covariance between genetic diversity and sexual reproduction at the neighborhood scale. We performed moving window analysis on local G/N (blue), reproductive effort (number of umbels produced / ramet height, RE, pink), reproductive success (number of fertile pods produced per umbel, RS, brown), and reproductive failure (proportion of pods aborted, RF, red). Circular window radii were 2m long, based on the inter-ramet distance in which there’s a 50% chance of being the same clone. (A) moving window rasters for Blandy Meadow, where white cells show no data (gray cells indicate low quantiles or zeros if zero-inflated), (B) Pearson’s correlation coefficients for raster cells were averaged across sites (points, lines show 95% confidence intervals). Points with confidence intervals that overlap 0 are in gray. Data shown for Blandy Meadow; all populations had similar patterns. See Fig S5 and Table S3 in the supplement.

### Does ramet sexual reproduction covary with genotype or spatial location?

Ramet reproductive effort, success, and failure were compressed into principal components (Fig 6A). The PCA explained ∼83% of fitness variation observed (Fig 6A); all three measures were strongly correlated with PC1 (correlations range −0.62 and −0.52, Table S4), while PC2 was driven primarily by a separation between reproductive success and reproductive failure.

**Figure 6.**
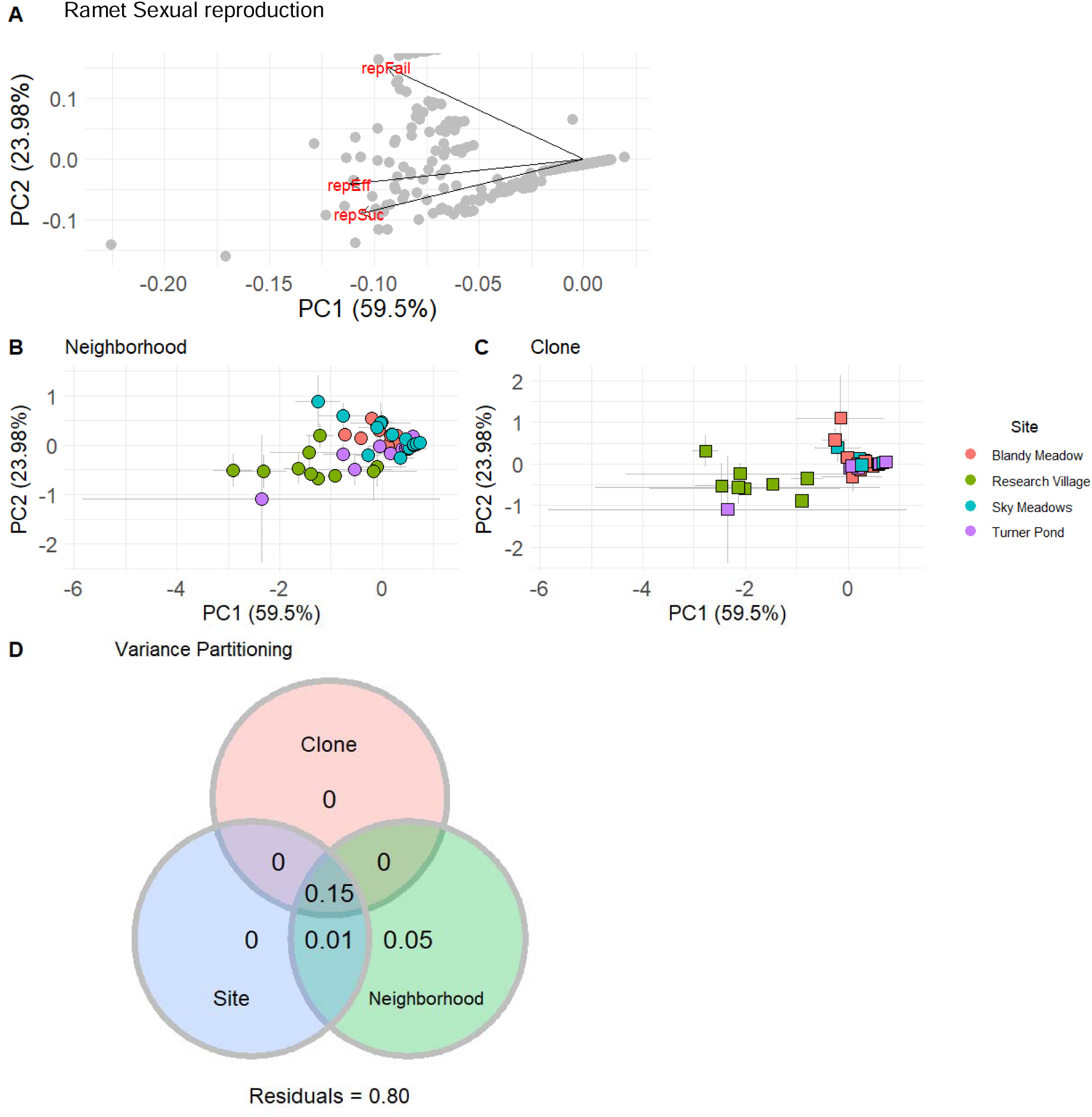
Ramet sexual reproduction is variable at multiple spatial scales and genotypes. (A) We compressed reproductive effort, success, and failure for milkweed ramets from 2023 using principal component analysis. (B,C) PCA centroids (averages) for neighborhood (B) -- where populations were divided equally into 16m^2^ neighborhoods -- and clone (C). Gray lines show 95% confidence intervals for PC1 and PC2. PERMANOVA and permutation tests for multivariate homogeneity (variance) were conducted for site, neighborhood, and clone. (D) Proportion of sexual reproduction explained by each predictor was determined by redundancy analysis and variance partitioning. Values represent the proportion of total variance in ramet sexual reproduction explained by each predictor uniquely or in combination with other variables. Residuals refers to residual variance.

When multivariate ramet sexual reproduction was examined, ramet phenotypes clustered by both space and genotype (Fig 6A, B). Sites were subdivided into 16 m^2^ neighborhoods to understand neighborhood-level effects that may not be captured with continuous distance. Site, neighborhood, and clone phenotypes occupied different sections of ordination space across all ramets observed in 2023 (Fig 6A, B). PERMANOVA revealed that site, neighborhood, and clone centroids were significantly different from one another (*P* < 0.001, Table S5), and the dispersion of multivariate homogeneity (or variance) was significantly different among sites, neighborhoods, clones (Table S5), *i.e.,* ramets in different clones, neighborhoods, and sites differ in their sexual reproduction and in their variation in sexual reproduction.

Because clones were unique to and spatially aggregated within populations, space and genotype are not independent. To better understand how each variable affects fitness uniquely, we conducted redundancy analysis (RDA) and quantified contribution of each variable to total variance explained with variance partitioning.

Though our predictors significantly modeled ramet sexual reprodcution, only 20% of variation in ramet sexual reproduction was described (*P* < 0.001, Table S6). Furthermore, variance partitioning revealed that only neighborhood explained unique variation, and even so, only 5% (Fig 6D). Only 1% of variance in sexual reproduction was described by the combination of site and neighborhood. Neighborhood, clone, and site in combination explained 15% of variance in phenotype. Clone identity, by contrast, did not significantly explain any variation in sexual reproduction among ramets (*P* > 0.4).

## Discussion

We observed similar demographic patterns across all four populations: approximately half of the genets we encountered were singletons and the remainder were a part of multi-ramet genets. The majority of multi-ramet genets were comprised of 5 or fewer ramets, but 9 of the 81 genets we observed were large (>20 ramets) and in three of our four sites the ramet population was dominated by only one or two clones. Clones were clustered within space, but within a clone, the ramets of small clones were dispersed and ramets of large clones were clustered. Both genetic diversity and reproductive success were spatially clustered; however, the spatial pattern had little effect on reproductive effort or success. Clone only explained variation in sexual reproduction among ramets in combination with site and environment.

The clonality rates that we estimate for *Asclepias syriaca* were similar to that estimated for the rare *Asclepias lanuginosa* (G/N = 0.11, Kim et al., 2014) and long-lived *Populus* species (G/N = 0.12, Dering et al., 2015). We found that clonal reproduction was much more prevalent in *A. syriaca* than had been previously described (Ricono et al., 2020): we found higher clonality (G/N of 0.1 in the current study vs. 0.8), a smaller proportion of singletons (51% vs. 86%), and more clustering of ramets within genets (69% probability that ramets within 1 meter were the same clone vs. 7%). These discrepancies stem from two differences between our studies. First, Ricono et al. (2020) used transect samples. Transect sampling can capture more spatial area, and thus more genetic variation, but will inevitably miss ramets within genets and bias estimates towards higher genetic diversity and inflate the number of singletons (Arnaud-Haond et al., 2007).

Second, Ricono et al., (2020) allowed for missing genetic data when determining multi-locus genotypes. Genetic mismatches between ramets were counted either if alleles were different or if allele data were missing in one or both ramets (Galpern et al., 2012), inflating the number of unique genotypes. Our sampling methods and genetic analysis provides a more robust and accurate picture of clonal structure and diversity in *A. syriaca*.

While most genets we encountered were small, in two of our populations, the single largest genet represented most of the ramet population (Research Village, 48%; Sky Meadows 85%) dominating the ramet population. In Blandy Meadow, two large clones dominated the ramet population (∼74% of ramets belonged one of two clones). In Turner Pond we found a different pattern with the largest clone comprising only 16% of the ramet population. The skewed demographic patterns we observed among these four populations have been reported in other taxa (Dering et al., 2015; Araki et al., 2022). In *Convallaria* clones (Araki et al., 2022), four dominant genotypes comprised 97% of the ramets studied, as we found in our three sites with a few dominant clones, albeit less skewed. However, the understory herb *Trillium recurvatum* balanced clonal and sexual reproduction almost equally (81 genets found in a population of 174 ramets, Mandel et al., 2019). However, in both studies, only a single population was sufficiently sampled. In our study, we observed substantial variation in genetic diversity and clone sizes among populations of the same clonal species, which highlights the importance of studying multiple populations to understand spatial dynamics of clonal plants.

Variation in genetic diversity and clone size among populations can be explained by population age, successional stage, or disturbance. Older populations may recruit more genets over time, increasing genotypic diversity through sexual reproduction and recombination (Wang et al., 2023). Immigration of new genotypes also play a key role; for example, in an *Erysimum* metapopulation with frequent recolonization, subpopulations showed varying degrees of genetic differentiation, though overall clonal diversity was stable (Honnay et al., 2008). In our study, Sky Meadows’ low genetic diversity may reflect its isolation within a forest matrix, limiting dispersal of seeds into the population. Disturbance may also play a role. A meta-analysis of clonal plants found that genotypic diversity tends to decrease with time since disturbance, as clonal growth replaces sexual reproduction (Silvertown, 2008). Though recent disturbance may favor sexual reproduction in most situations (Silvertown, 2008), under dispersal limitation, disturbance could further decrease competitive pressure and allow one genotype to spread clonally and dominate. Some species, particularly seagrasses, were found to rely on clonal reproduction following disturbance (Silvertown, 2008).

We found evidence that ramet growth patterns shift as clones grow larger. Small genets spread ramets more widely, likely to increase resource access and limit self-competition, a behavior seen in *Hydrocotyl vulgaris* (Zhang et al., 2022). In contrast, larger genets grow more clustered, possibly due to the higher carbon cost of spreading further, or because large genets can grow multiple ramets in favorable microsites (adaptive foraging, Fischer & Van Kleunen, 2001; Roiloa & Retuerto, 2006; Weiser et al., 2016). This shift in growth pattern could reflect ontological traits like adventitious root elongation or root bud production, as genet size, area, and ramet density were consistent across sites.

The mechanisms and drivers of bud production on clonal growth organs may explain patterns of ramet distribution within genets. In *A. syriaca*, Polowick & Raju (1982), found higher bud production near the base of ramets, especially just after seedling establishment. These data, however, suggest the opposite growth pattern than we observed in small genets, though we do not know the age of the small genets in our study. Age and size may be separate drivers of clonal growth. Diversion of resources towards buds may also play a role. Research on milkweed invasions found that ramets distribute herbicide to distant buds more than those near the base (Waldecker & Wyse 1985; Backacsy & Bagi, 2020), suggesting that vascular systems may promote circulating resources further away. Additionally, buds in milkweed may develop facultatively, responding to resource availability, or obligately as part of overwintering (Klimesova & Martinkova, 2004). The production and dormancy of buds, though understudied, likely plays a key role in the distribution of genetic variation within populations (Herben & Klimesova, 2020). Understanding bud production dynamics may provide insight into clonality, spatial structure and resource allocation in perennial plants.

In our study, there was no relationship between local genetic diversity and reproductive success or failure. Our moving window analyses showed that clonal diversity was high within certain patches of each population, creating pockets of genetic variability that are sufficient to support higher pollination success (Wyatt & Broyles, 1994; Dupont et al., 2014). We did not observe widespread reproductive failure, suggesting that genotypic diversity was sufficient to promote outcrossing for successful seed production. Outcrossing among populations may also be more common through pollinator foraging behavior. *Bombus* is consistently found to be an effective pollinator in milkweeds (Stoepler et al., 2012; Gustafson et al., 2023) that can travel nearly 1km when foraging (Knight et al., 2005), sufficient for visiting genetically-variable pollen donors and connect populations with low genetic diversity. In our study, ramets with high reproductive success also experienced higher reproductive failure through increased self-pollination events. However, the cost of ovule abortion may be low if cloning allows genotypes to persist and attempt sexual reproduction in better years (Vallejo-Marin et al., 2010).

Clone, neighborhood, and site simultaneously explained 15% of variation in ramet sexual reproduction. However, neighborhood was the only component that uniquely explained variation. Milkweed is highly plastic, as shown in recent study in the genetic and environmental basis of common milkweed size and herbivore defense phenotypes (Potts & Hunter, 2021).

Maternal lines grown in a field and greenhouse common garden showed no differences in cardenolide concentration, but some genetic basis for height and leaf production (Potts & Hunter, 2021). In our study, similarities among ramets within genets were primarily due to their spatial proximity, and not their genotype nor local genetic variation. Similar to our results, Potts & Hunter (2021) found that common garden location and experimental block (i.e. sublocation) affected ramet phenotype, showing a scale-dependent response in ramet phenotype. Interestingly, the lack of variance explained by site indicates that site-level differences in our study were driven by the cumulative effect of variation among neighborhoods. One caveat to this analysis is that we are only representing genotypes from a single region of milkweed’s range, and ecotypes are known to vary in their biomass allocation patterns and likely fitness (Woods et al., 2012; De La Mater et al., 2018).

For clonal plants, genotype may better predict other aspects of clonal reproduction, not the sexual reproduction of ramets. For root sprouters, below-ground biomass (rootstock), bud production, and degree of physiological integration could be critical measures of fitness (Herben & Klimesova, 2020). Genotypes across the range of milkweed showed variation in aboveground biomass allocation (Woods et al., 2012), and this may extend to root characteristics as well.

Additionally, variation in ramet phenotype due to microsites differences may be masked if genets can distribute resources among ramets. For example, in a rhizomatous grass, physiological integration increased biomass production, particularly in response to grazing and interspecific competition (Evans et al., 2023). Future work should seek to understand ecologically-relevant genet phenotypes in milkweed, and its variation across spatially heterogeneous environments.

### Conclusion

High rates of clonality in common milkweed populations in Northern Virginia create a skewed demographic structure of ramets and genets, where few large clones comprised up to 86% of the ramet population. Clonal growth patterns showed that ramets of the same genotype were more likely to interact but may undergo dispersive growth to avoid competition with self. Furthermore, our research shows that clonal plant populations can showcase high sexual reproduction even with low genetic variability. In addition, neither local genetic variation nor genotype predicted sexual reproduction of ramets, but rather neighborhood. Our research shows that clonality can structure demographic and competitive dynamics in clonal plants while maintaining high sexual reproduction.

## Acknowledgements

This work was supported by the University of Virginia Blandy Experimental Farm Graduate Student Research Fellowship, The Garden Club of America, a Crouch Memorial Fellowship from William & Mary, and the Virginia Native Plant Society. Special thanks to Dave Carr, Kyle Haynes, and Blandy staff for field coordination. Phoebe Williams of William & Mary, Jan MacDowell of the Virginia Institute of Marine Science, and Linda Cote of Cornell Biotechnology Institute provided guidance and support with molecular methods and genotyping.

M. Drew LaMar advised us during data analysis. Many thanks to William & Mary field and lab assistants Ivan Munkres, Casey Hensen, Geneva Waynick, Josh Mutterperl, Mia Perry, Katie Barlow, Grace Gulbankian, Helena Huber, Owen George, Anika Kalluri, Bela Rein, Grace Parker, and Cameron Morris.

## Author contributions

H.M.M., H.J.D., and J.R.P. designed this research project. H.M.M. and H.J.D. collected the field data, H.M.M. and J.R.P. developed molecular methods, and H.M.M. processed genetic samples. H.M.M. performed the analysis and wrote the manuscript with contributions from all authors. All authors contributed to the review of the research and writing.

## Supporting Information

### Methods S1. Sampling simulation

We determined the best sampling scheme for a milkweed population to best capture the spatial genetic relationship of milkweed clones as well as ramet demography (number of genets and their size). We simulated x,y coordinates for 50 populations of 450 ramets in a 40m x 40m square. Each “ramet” was assigned a genet ID following a phalanx (clustered), guerrilla (random), or intermediate growth pattern (clustered with 50% of ramets reassigned randomly).

We performed a factorial simulation modifying the growth form and the ratio of genotypes to population size (G/N, at 0.25 and 0.75), producing 6 population types with a sample size of 50. In all populations, 10% of ramets were randomly reassigned to genets to add noise. From these populations, ramets were sampled following two methods– simple random sampling or random clustered sampling (nearest neighbors)– at different proportions of the population (90-20% sampled). We then evaluated how sampling scheme, and sampling proportion affected three estimates of clonal demography relative to that of the true population estimates (100% sampled): G/N, the clone size distribution, and the spatial genetic relationship of clones. The spatial genetic relationship was classified as the distance coefficient and intercept of a binomial GLM of the probability of being the same clone as a function of inter-ramet distance. We quantified the proportion of models (out of 50) that fell within the 95% confidence interval generated from a population census. The genet size distribution was sensitive to sampling effort. We found that sampling 75% or more of the population under phalanx and intermediate scenarios with few genets (G/N = 0.25) enabled us to capture both the spatial genetic relationship and clonal demography of the population that fell within the 95% confidence interval of the true population estimate of each.

**Figure S1.**
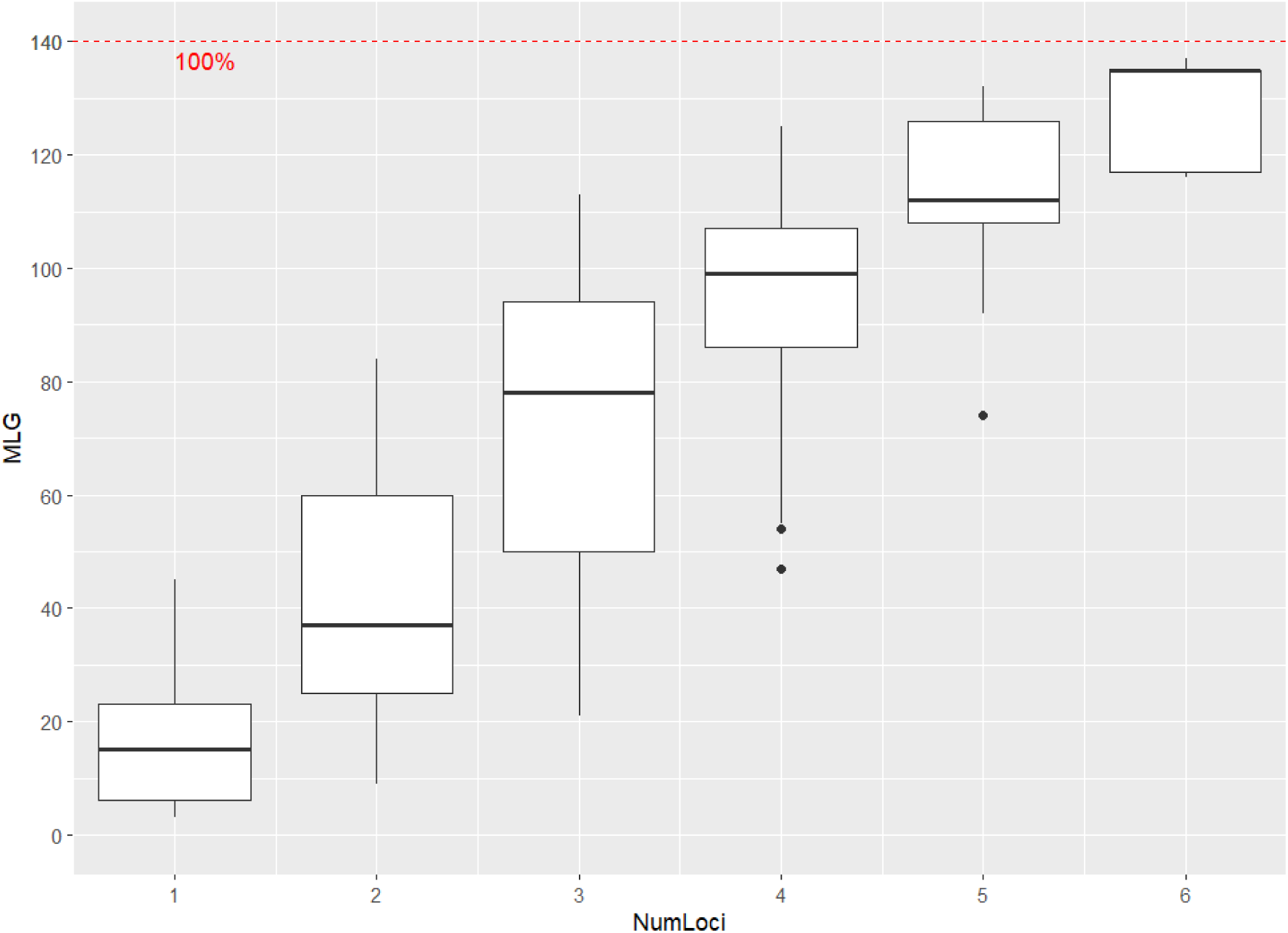
Genotype accumulation curve for milkweed microsatellites. 7 microsatellite genotypes published in Kabat et al. (2010) were amplified in 1629 milkweed ramets. Genotype accumulation curve was generated using the *genotype_curve* function in *adegenet* with 1000 iterations.

**Figure S2.**
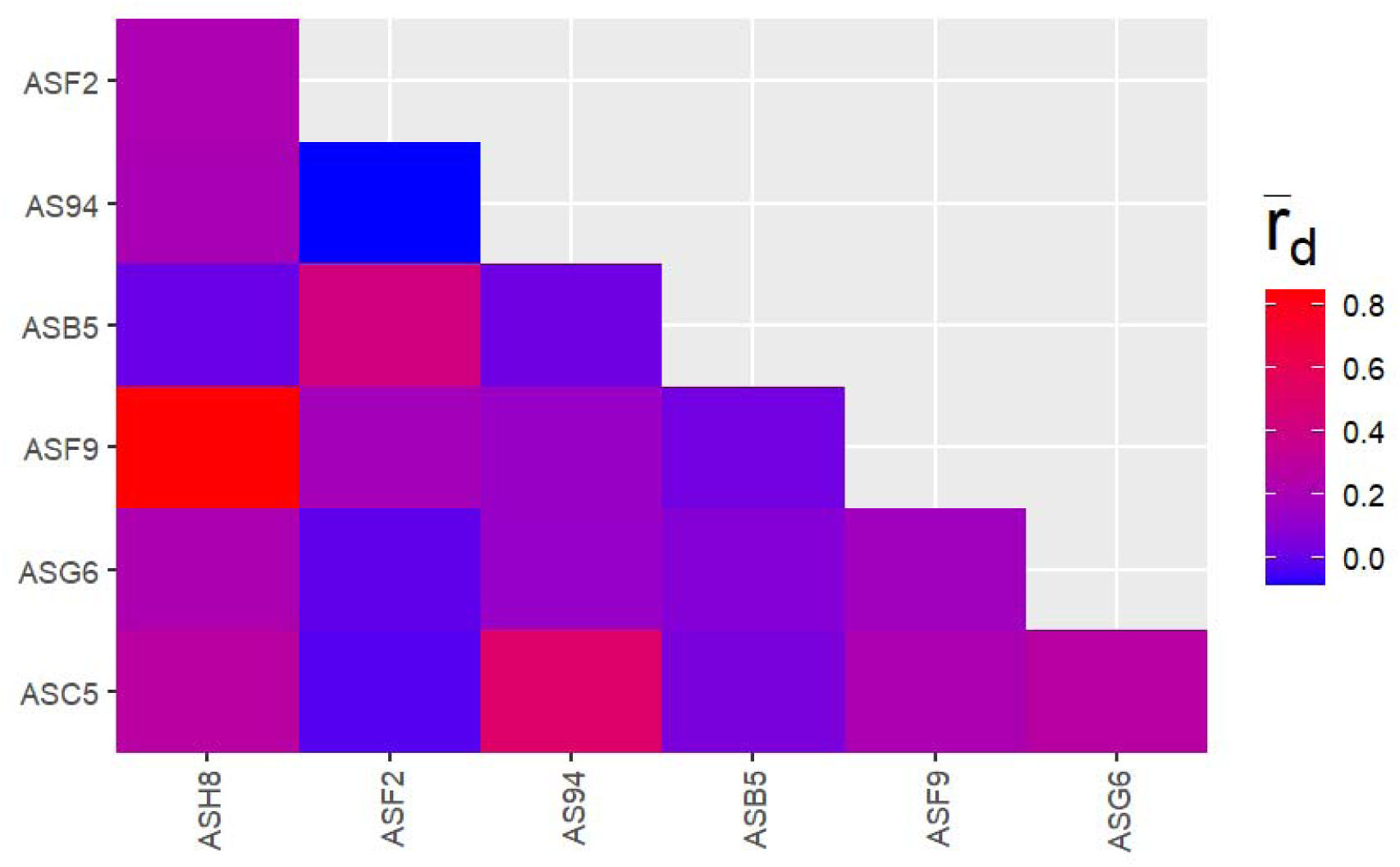
Pairwise standardized index of association (rd) among 7 microsatellite loci. Milkweed microsatellites (Kabat et al. 2010) were amplified from four populations of common milkweed in 2023. Pairwise rd was calculated using the pair.ia function in poppr.

**Figure S3.**
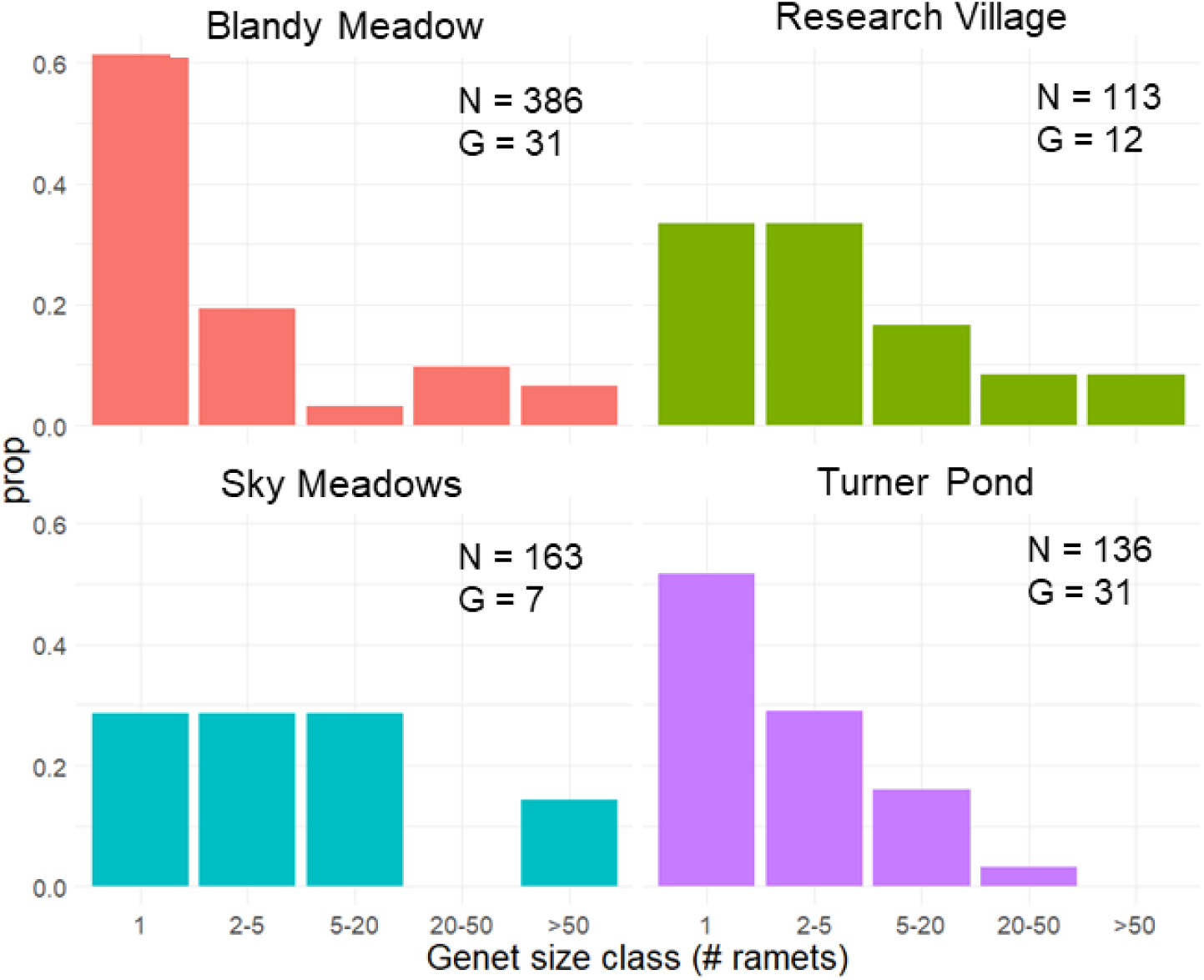
In four populations, most milkweed genets were small. Bars show the proportion of genets in different size classes (number of ramets per genet) for four population in Northern Virginia in 2023. Clones were identified from multi-locus genotypes at 7 microsatellite loci published in Kabat et al. (2010).

**Figure S4.**
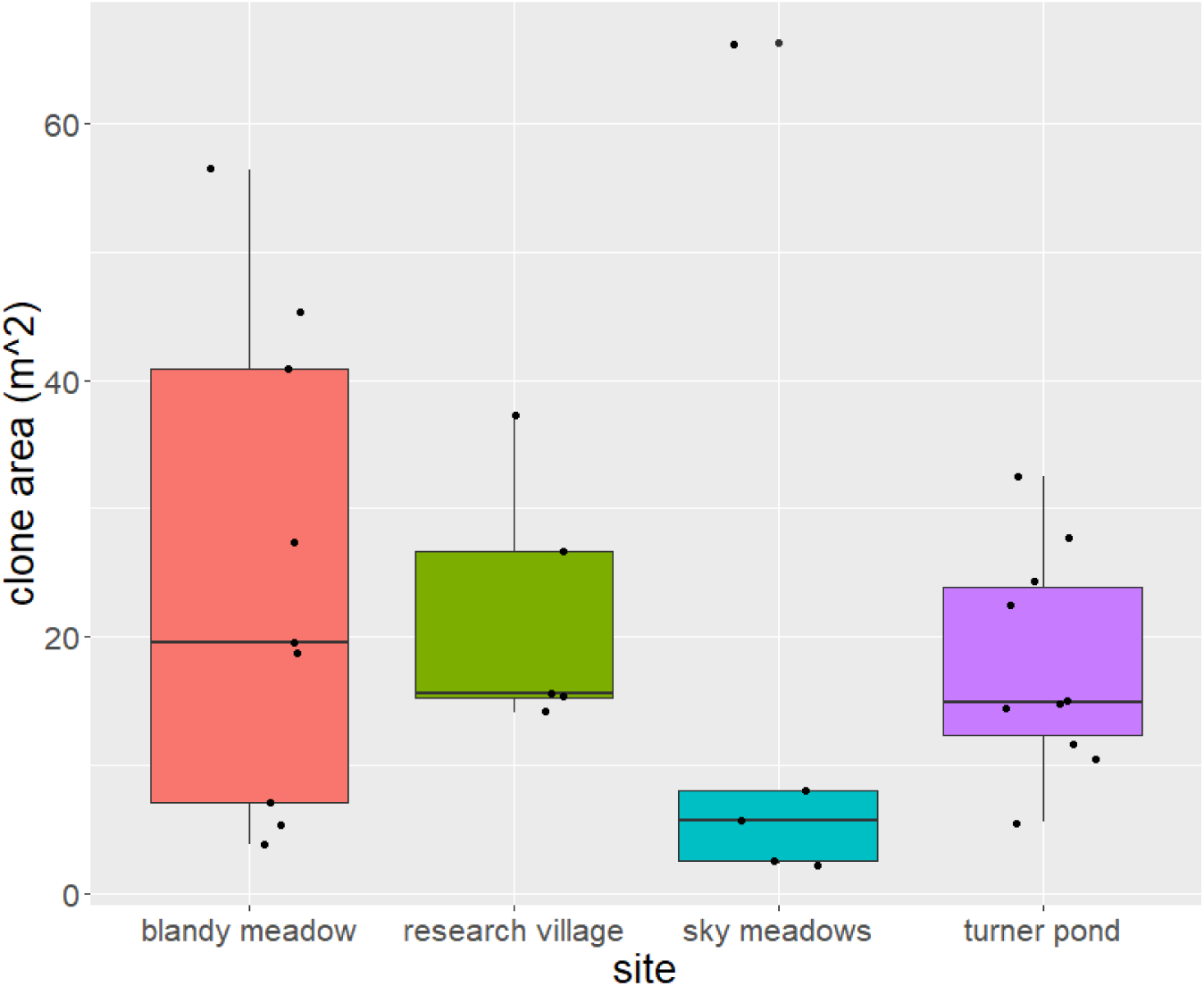
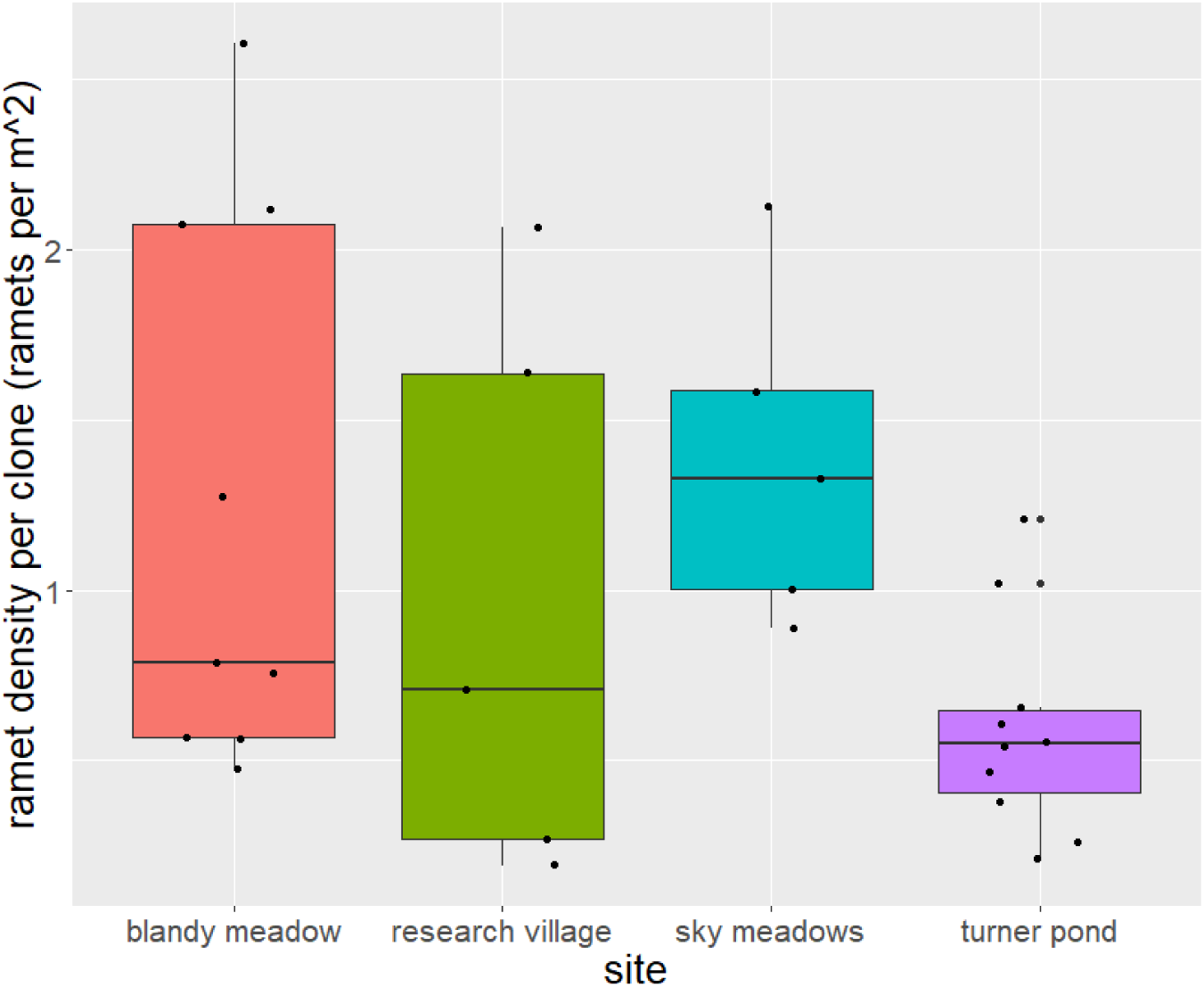
Clone area and ramet density variable among sites in 2023. For each multi-ramet clone (>3 ramets), we calculated the minimum convex polygon surrounding all ramets within the clone (clone area) using the mcp() function in the adehabitatHR package, then divided the number of ramets within that clone by its area (ramet density).

**Figure S5.**
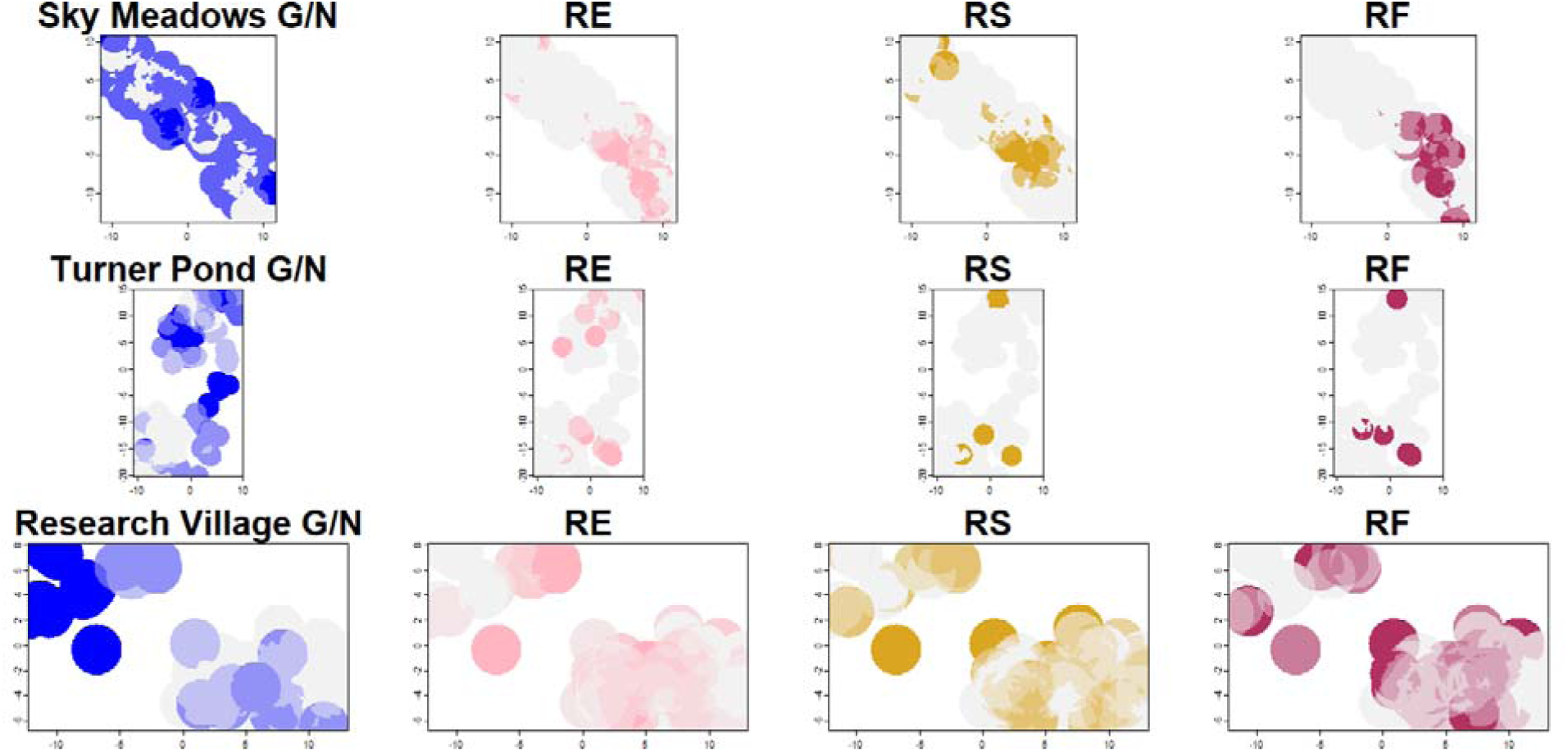
Spatial structure of local genetic diversity and reproduction in common milkweed sites. We performed a moving window analysis on local clonal diversity (G/N, blue), reproductive effort (number of umbels produced / ramet height, pink), reproductive success (number of fertile pods produced per umbel, brown), and reproductive failure (proportion of pods aborted, red) using the focal() function in the terra package in R. Circular window radii were 2m long, based on the inter-ramet distance in which there’s a 50% chance of being the same clone. Data represent quantiles <0.1, 0.25, 0.5, and >0.75 for raster values. White cells have no data.

**Figure S6.**
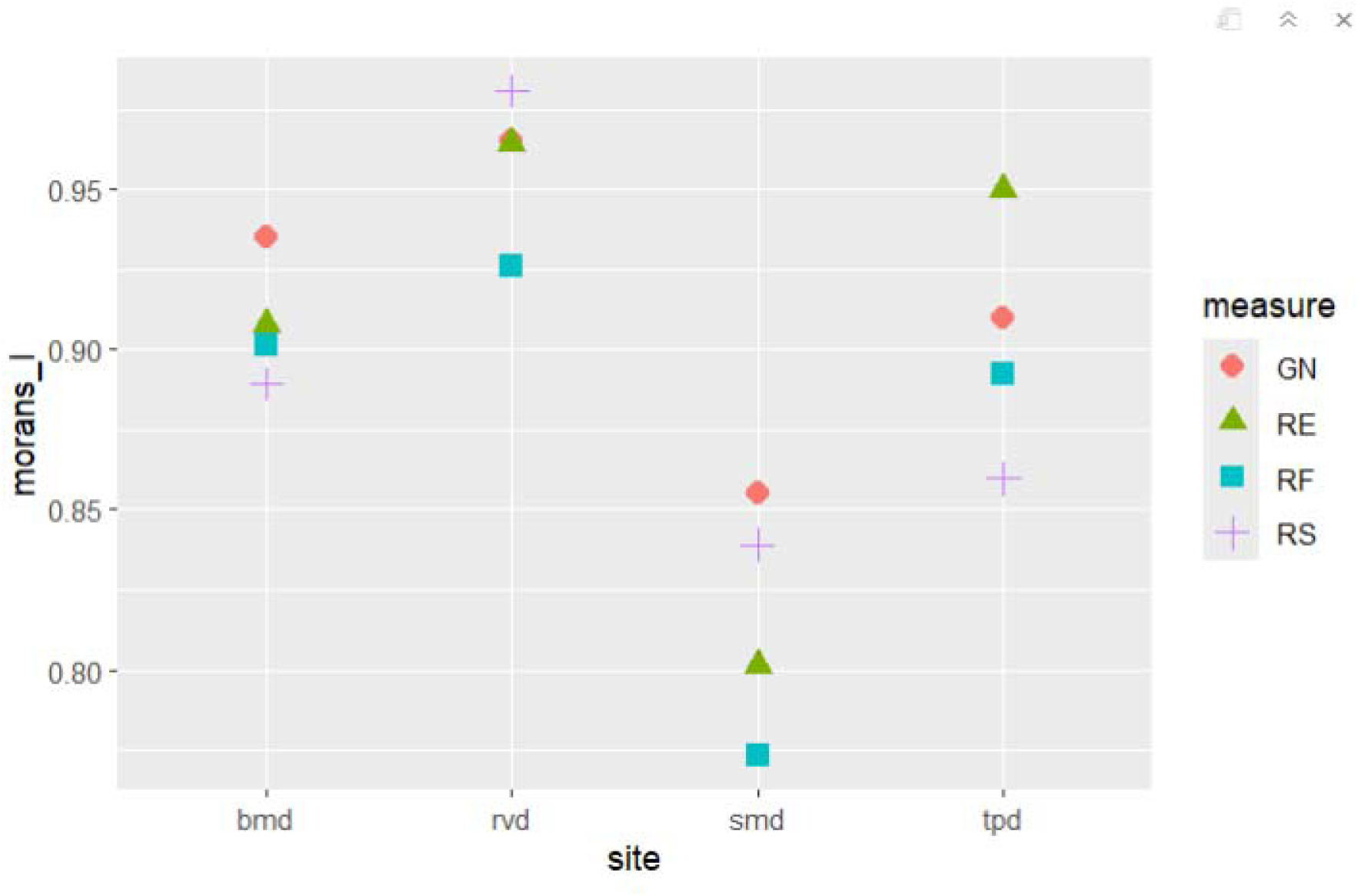
Spatial autocorrelation of local genetic diversity and sexual reproduction is highly clustered across sites. Points show global Moran’s I values calculated from rasters generated from moving window analysis (see text for details). N_BM_=165511, N_TP_ = 173963, N_SM_ =114960, N_RV_ =78698 cells.

**Table S1.** Genetic distance is correlated with spatial distance across time in two milkweed populations. Spatial and genetic data from two milkweed populations were collected from 2021 to 2023. Genetic distance was calculated as the Bray Curtis distance between ramets for alleles at 7 microsatellite loci (Kabat et al. 2010). Phenotypic distance was measured as Euclidean distance of ramet height, herbivory, umbel production, and pod production. All p-values < 0.001.

| Site | Mantel Correlation<br>Coefficient<br>(Distance x Genotype) |
| --- | --- |
| Blandy Meadow | 0.41 |
| Turner Pond | 0.36 |
| Sky Meadows | 0.20 |
| Research Village | 0.39 |

**Table S2.** Spatial-genetic relationships of common milkweed populations. Binomial generalized linear models were fit to ramet-ramet comparisons of genetic similarity (same or different clones) and physical distance for each population and year. All parameters are significant (p<0.001).

| Site | Intercept $\beta \pm$<br>SE | Distance $\beta \pm$ SE | Residual<br>deviance | Null<br>deviance | Pseudo-$R^2$ |
| --- | --- | --- | --- | --- | --- |
| Blandy Meadow | $0.387 \pm 0.02$ | $-0.204 \pm 0.002$ | 70792 | 79655 | 0.11 |
| Turner Pond | $0.730 \pm 0.084$ | $-0.405 \pm 0.016$ | 3442.8 | 5493.3 | 0.37 |
| Research Village | $1.79 \pm 0.07$ | $-0.429 \pm 0.013$ | 5079.3 | 7701.4 | 0.34 |
| Sky<br>Meadows | $1.80 \pm 0.041$ | $-0.060 \pm 0.003$ | 14382 | 14817 | 0.03 |

**Table S3.** Covariance of phenotypes on landscapes at the neighborhood scale. Pearson’s correlation coefficients were calculated for cells in moving window rasters generated for local G/N (genotypes per ramets sampled), reproductive effort (number of umbels produced / ramet height, RE), reproductive success (number of fertile pods produced per umbel, RS), and reproductive failure (proportion of pods that were aborted, RF). Circular window radii were 2m long, based on the inter-ramet distance in which there’s a 50% chance of being the same clone. Bold values represent significant coefficients (p<0.001).

| Pop |  | G/N | RE | RS | RF |
| --- | --- | --- | --- | --- | --- |
| Blandy<br>Meadow | G/N |  |  |  |  |
|  | RE | 0.124918 |  |  |  |
|  | RS | 0.170281 | 0.793481 |  |  |
|  | RF | 0.087504 | 0.809631 | 0.7925 |  |
|  |  | G/N | RE | RS | RF |
| Turner<br>Pond | G/N |  |  |  |  |
|  | RE | 0.2407 |  |  |  |
|  | RS | -0.21515 | 0.096929 |  |  |
|  | RF | -0.21908 | 0.100437 | 0.750308 |  |
|  |  | G/N | RE | RS | RF |
| Research<br>Village | G/N |  |  |  |  |
|  | RE | 0.059436 |  |  |  |
|  | RS | 0.404488 | 0.525897 |  |  |
|  | RF | 0.06646 | 0.397131 | 0.34073 |  |
|  |  | G/N | RE | RS | RF |
| Sky<br>Meadow<br>s | G/N |  |  |  |  |
|  | RE | -0.22557 |  |  |  |
|  | RS | -0.13576 | 0.652405 |  |  |
|  | RF | -0.21103 | 0.741375 | 0.527238 |  |

N<sub>BM</sub>=165511, N<sub>TP</sub> = 173963, N<sub>SM</sub> =114960, N<sub>RV</sub> =78698

**Table S4.** Correlations among ramet phenotypes and principal component axes. Principal component analysis was performed on reproductive effort, success, and failure using data from ramets in four population in 2023.

|  | PC1 | PC2 | PC3 |
| --- | --- | --- | --- |
| RE | -0.6180877 | -0.2296786 | -0.7518080 |
| RS | -0.5903422 | -0.4959108 | 0.6368426 |
| RF | -0.5190988 | 0.8374486 | 0.1709274 |

**Table S5.**
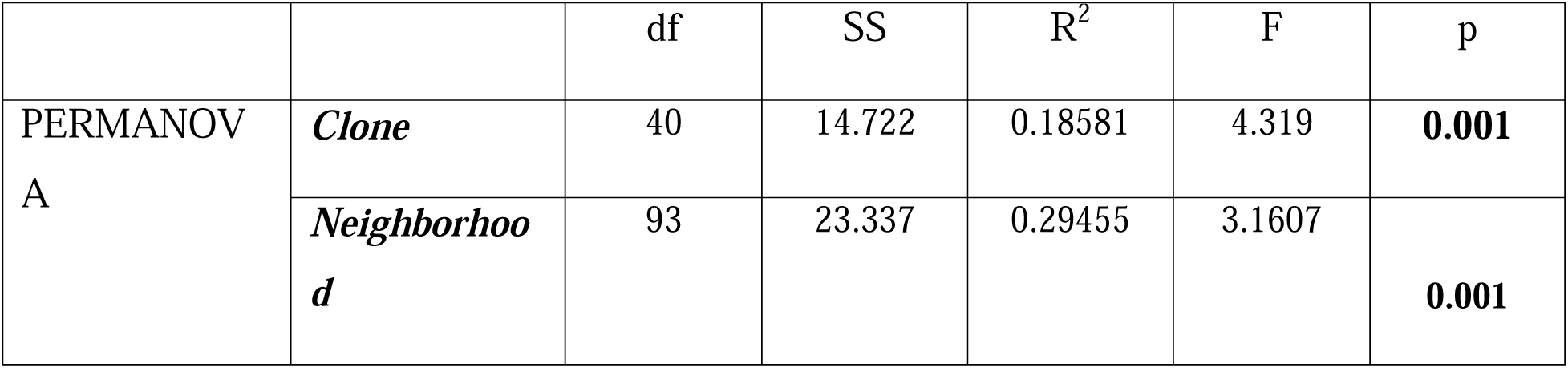
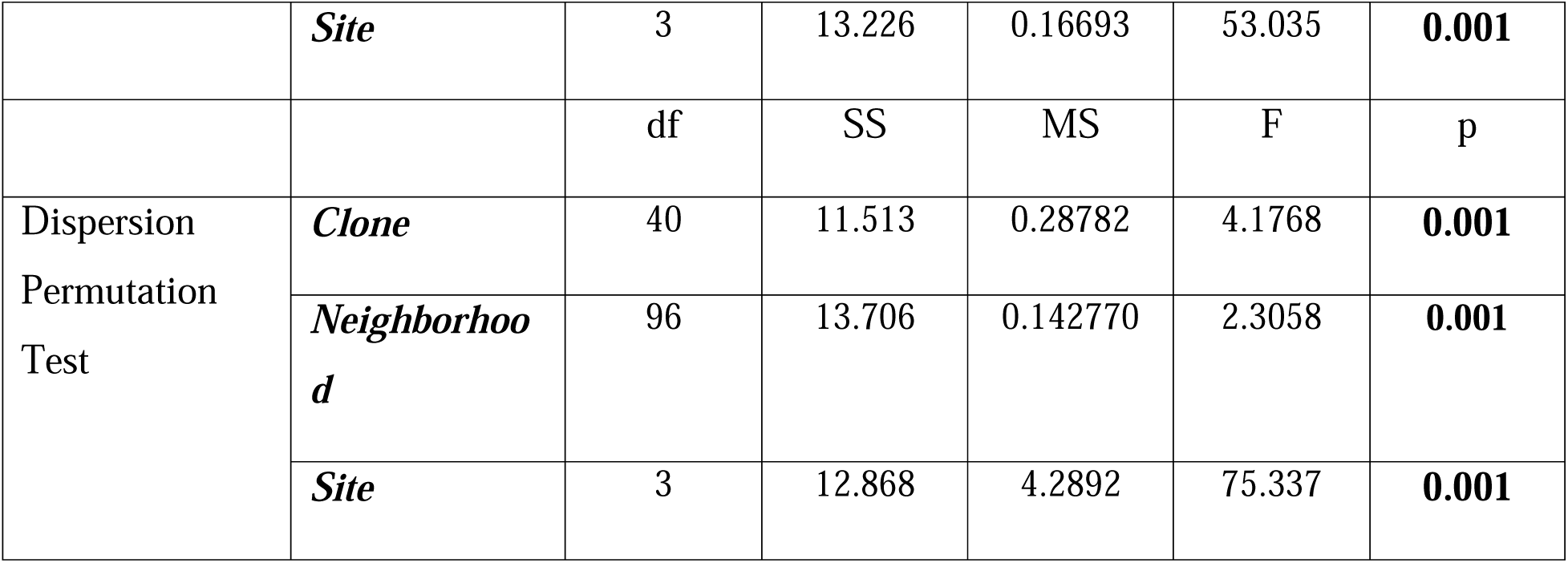
Ramet sexual reproduction clusters by neighborhood, site, and clone in all sites in 2023. Phenotypes include height, reproductive effort, and reproductive success. Each population was subdivided into 16 neighborhoods. PERMANOVA and permutation tests for multivariate homogeneity of groups was performed using the vegan package using data from all ramets (n = 797).

**Table S6.**
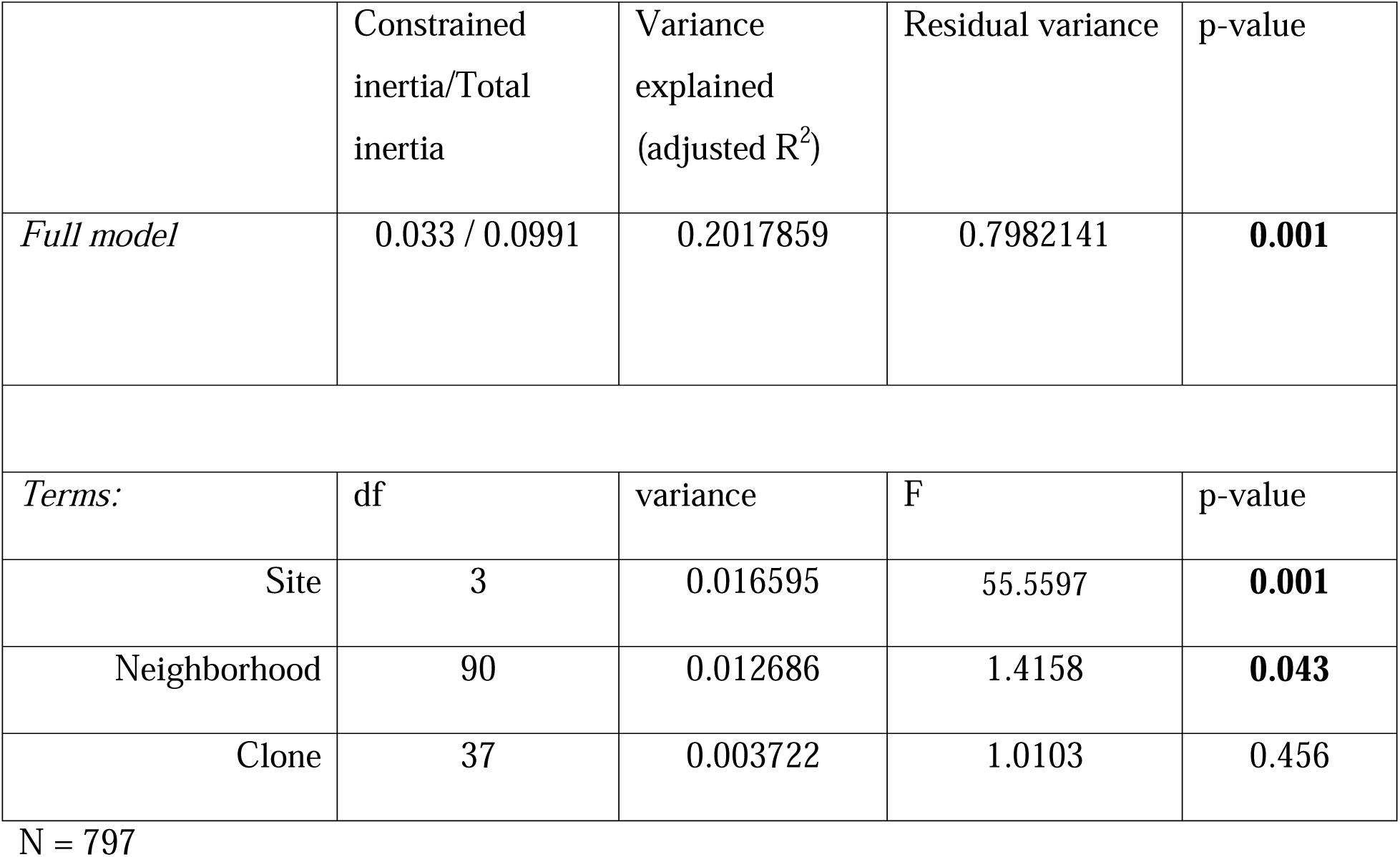
Summary of redundancy analysis of ramet phenotype and genotype, quadrant, and site. Redundancy analysis was performed on ramets from 2023 with ramet height, herbivory, reproductive effort, and reproductive success as responses and neighborhood, site, and clone ID as predictors. Significance was evaluated with ANOVA using the anova.cca() function in vegan.

